# Photoreactive hydrogel stiffness influences volumetric muscle loss repair

**DOI:** 10.1101/2021.06.19.449065

**Authors:** Ivan M. Basurto, Juliana A. Passipieri, Gregg M. Gardner, Kathryn K. Smith, Austin R. Amacher, Audrey I. Hansrisuk, George J. Christ, Steven R. Caliari

## Abstract

Volumetric muscle loss (VML) injuries are characterized by permanent loss of muscle mass, structure, and function. Hydrogel biomaterials provide an attractive platform for skeletal muscle tissue engineering due to the ability to easily modulate their biophysical and biochemical properties to match a range of tissue characteristics. In this work we successfully developed a mechanically tunable hyaluronic acid (HA) hydrogel system to investigate the influence of hydrogel stiffness on VML repair. HA was functionalized with photoreactive norbornene groups to create hydrogel networks that rapidly crosslink via thiol-ene click chemistry with tailored mechanics. Mechanical properties were controlled by modulating the amount of matrix metalloproteinase (MMP)-degradable peptide crosslinker to produce hydrogels with increasing elastic moduli of 1.1 ± 0.002, 3.0 ± 0.002, and 10.6 ± 0.006 kPa mimicking a relevant range of developing and mature muscle stiffnesses. Functional muscle recovery was assessed following implantation of the HA hydrogels by *in situ* photopolymerization into rat latissimus dorsi (LD) VML defects at 12 and 24 weeks post-injury. After 12 weeks, muscles treated with medium stiffness (3.0 kPa) hydrogels produced maximum isometric forces most similar to contralateral healthy LD muscles. This trend persisted at 24 weeks post-injury, suggestive of sustained functional recovery. Histological analysis revealed a significantly larger zone of regeneration with more *de novo* muscle fibers following implantation of medium stiffness hydrogels in VML-injured muscles compared to other experimental groups. Lower (low and medium) stiffness hydrogels also appeared to attenuate the chronic inflammatory response characteristic of VML injuries, displaying similar levels of macrophage infiltration and polarization to healthy muscle. Together these findings illustrate the importance of hydrogel mechanical properties in supporting functional repair of VML injuries.

## 1. Introduction

Skeletal muscle is a dynamic tissue capable of adapting and remodeling in response to external stimuli to maintain homeostasis in the presence of tissue damage^1^. However, in the case of certain congenital and acquired diseases or traumatic injuries, loss of muscle fibers and surrounding tissue exceeds the endogenous regenerative capacity and results in a permanent reduction of muscle mass and function, as well as poor cosmetic outcomes. Injuries of this nature have been termed volumetric muscle loss (VML)^2,3^. While medical advances have decreased fatalities in recent military conflicts^4^, traumatic injuries persist. In fact, damage to the musculoskeletal system constitutes the majority of injuries sustained by current war fighters^2^ with 53% of combat-related extremity injuries resulting in open skin wounds that directly affect soft tissues^5^. VML injuries also affect the civilian population where skeletal muscle damage results from various myopathies, as well as surgical loss, traumatic injuries (e.g., car accidents, gunshot wounds), and compound fractures^6^. Current approaches to treating severe muscle injuries include the autologous transfer of healthy skeletal muscle from a donor site to the area of injury followed by extensive physical therapy^5,7^. However, this treatment method often results in poor levels of functional improvement as well as associated donor site morbidity^7–9^. Therefore, there is a clear need for therapies that more fully restore muscle form and function.

In an effort to address this unmet medical need, tissue engineering approaches have emerged as a promising therapeutic platform^10–13^. Hydrogel biomaterials provide an attractive alternative to tissue and/or extracellular matrix (ECM)-based treatments for VML injuries owing to the variety of different material systems and chemistries available that can be tailored for specific skeletal muscle tissue engineering applications^14–16^. In the case of VML injuries, the use of photopolymerizable hydrogels holds promise due to their rapid gelation kinetics, facile delivery, conformability to the wound site^17,18^, and high level of chemical tunability^19^. Hydrogel properties can be easily modulated to recapitulate salient biophysical features including soft tissue mechanics^20,21^. Due to the ability to recreate distinct tissue characteristics, hydrogels have been used extensively to study the complex interplay between cells and their microenvironment^22,23^.

Prior work has shown that cells can sense ECM biomechanics by translating environmental cues into intracellular signals by a process known as mechanotransduction^24^. Mechanotransduction can regulate many cell behaviors including spreading, migration, and stem cell differentiation^25–28^. Pioneering work in the field of mechanobiology showed that mesenchymal stromal cells (MSCs) cultured atop hydrogels with varying stiffness preferentially differentiated toward brain, muscle, or bone lineages with increasing substrate elasticity^27^. More recent studies have built on these findings and shown that hydrogels with Young’s moduli (stiffnesses) of 10-12 kPa, a stiffness matching atomic force microscopy (AFM) measurements of muscle tissue stiffness^29^ and previously reported to promote MSC myogenic differentiation *in vitro*^27,29^, supported proliferation of skeletal muscle stem cells (satellite cells) *in vitro* and improved engraftment efficiency when transplanted *in vivo*^30^. While these findings have been informative to the study of how cells interact with their local microenvironment, they have not been thoroughly explored in the context of tissue engineering and wound healing.

Currently, natural VML wound healing is characterized by an elevated and sustained immune response that results in fibrotic collagen deposition, ECM stiffening, and minimal restoration of muscle mass and function^31^. This response is remarkably different than fetal and neonatal wound healing where tissues experience minimal inflammation and reduced scarring^32–34^. One potential mediator of this disparate tissue response is the gradual transition of the skeletal muscle niche from a more compliant biomechanical state to a stiffer, more elastic environment^35–37^. Although measurements of embryonic muscle mechanics are difficult to make, recent AFM analysis of 8-day old embryonic chick midgut muscularis reported elasticity on the order of 0.5 kPa^38^. Myofibers isolated from 3-month-old rats had stiffnesses of around 0.5-2 kPa^39^. Additionally, skeletal muscle ECM has been shown to stiffen with age and is characterized by augmented collagen deposition and limited muscle progenitor cell proliferation^40^. These developmental changes in the skeletal muscle microenvironment, specifically the increased stiffness, could result in reduced regenerative outcomes. Therefore, we hypothesized that a compliant hydrogel system with a lower elastic modulus than adult muscle (10-12 kPa) would result in improved functional tissue recovery while inhibiting fibrosis. Here we describe our results with a hydrogel platform with tunable mechanical properties to investigate the influence of hydrogel stiffness on functional VML repair.

## 2. Materials and Methods

### 2.1 NorHA synthesis

Hyaluronic acid (HA) was functionalized with photoreactive norbornene groups as previously described19. Briefly, sodium hyaluronate (Lifecore, 62 kDa) was first converted to hyaluronic acid *tert*-butyl ammonium salt (HA-TBA) using Dowex 50W proton exchange resin. HA-TBA was subsequently filtered, titrated to a pH of 7.05, frozen, and lyophilized. Norbornene groups were then added by a reaction with 5-norbornene-2-methylamine and benzotriazole-1-yloxytris-(dimethylamino) (BOP) in dimethylsulfoxide (DMSO). The reaction was conducted at 25°C for 2 hours prior to being quenched with cold deionized (DI) water and transferred to dialysis tubing. Finally, NorHA was dialyzed for 5 days, filtered, dialyzed for an additional 5 days, frozen, and lyophilized. The degree of HA modification with norbornene was calculated at 22% as determined via proton nuclear magnetic resonance (^1^H NMR) (Fig. S1).

### 2.2 Degradable peptide synthesis

Solid-state peptide synthesis was conducted on a Liberty Blue automated microwave-assisted peptide synthesizer. A matrix metalloproteinase (MMP)-degradable peptide (GCNSVPMSMRGGNCG, degradation occurs between the serine and methionine residues) that is susceptible to degradation by MMP-1 and MMP-2^41^ was synthesized on a Rink Amide MBHA high-loaded (0.78 mmol/g) resin using solid-supported Fmoc-protected peptide synthesis. The resin was swelled with 20% (v/v) piperidine in dimethylformamide (DMF) and the amino acids were sequentially added from C to N-terminus. The resulting peptide was collected and cleaved in 92.5% trifluoroacetic acid, 2.5% triisopropylsilane, 2.5% (2,2-(ethylenedioxy)diethanethiol), and 2.5% H2O for 2 hours and filtered to separate the resin. The peptide was precipitated in cold ether, dried overnight, resuspended in water, frozen, and lyophilized. Synthesis was confirmed via matrix-assisted laser desorption/ionization (MALDI) (**Fig. S2**).

### 2.3 NorHA hydrogel fabrication

NorHA hydrogels were fabricated via an ultraviolet (UV) light-mediated thiol-ene click reaction between norbornene groups and terminal thiols. Hydrogel precursor solution was prepared by incorporating thiolated RGD peptide (GCGYGRGDSPG, 1 mM, Genscript) to promote integrin-mediated cell adhesion and lithium acylphosphinate (LAP, 1 mM) to serve as the photoinitiator into a 6 wt% NorHA stock solution. Hydrogel mechanical properties were then modulated by adjusting the concentration of dithiol-containing MMP-degradable peptides in the precursor solution and photopolymerized using 365 nm UV light at an intensity of 7 mW cm-2.

### 2.4 Mechanical characterization

NorHA hydrogel mechanical properties were assessed using an Anton Paar MCR 302 oscillatory shear rheometer. Gelation kinetics and storage moduli were quantified using a cone-plate geometry (25 mm diameter, 0.5°, 25 μm gap) via oscillatory time sweeps (1 Hz, 1% strain). To facilitate *in situ* gelation NorHA precursor solutions were irradiated for 2 minutes with UV light (365 nm) at an intensity of 7 mW cm^−2^. Hydrogel elastic moduli were calculated as 3 times the storage modulus assuming a Poisson’s ratio of 0.520. All experimental groups were tested in triplicate.

### 2.5 Animal care

This study was conducted in compliance with the Animal Welfare Act, the Implementing Animal Welfare Regulations, and in accordance with the principles of the Guide for the Care and Use of Laboratory Animals. The University of Virginia Animal Care and Use Committee approved all animal procedures. A total of 65 male Lewis Rats (Charles River Laboratories) age matched to 12 weeks weighing 312.7 ± 24.9 g were purchased and individually housed in a vivarium accredited by the American Association for the Accreditation of Laboratory Animal Care and provided with food and water *ad libitum*.

### 2.6 Surgical procedures

The VML model was created by resecting approximately 13% of the total muscle mass from the latissimus dorsi (LD) of male Lewis rats as previously described^3,42,43^. Briefly, animals were anesthetized via isoflurane and the surgical site was aseptically prepared by repeated washes with alcohol and iodine. A longitudinal skin incision was made along the midline of the back and the fascia was separated to expose the underlying muscles. Next, the trapezius muscle was cut and lifted to access the underlying LD muscle. Once the LD was exposed, a 1.5 × 1.2 cm rectangle was drawn along the lateral edge of the muscle using surgical markers to represent the dimensions of the defect. The VML injury was then created by resecting a rectangular area (thickness: 1-2 mm) using surgical scissors, which corresponded to 148.6 ± 10.4 mg of tissue.

After creation of the defect, animals were randomly assigned to one of the following experimental groups: no repair (NR; *n* = 17), low (1 kPa, *n* = 16), medium (3 kPa, *n* = 16), and high (10 kPa, *n* = 16) stiffness hydrogels. A custom designed, pliable rectangular mold with the same dimensions as the VML injury was printed at the University of Virginia Mechanical and Aerospace Engineering (MAE) rapid prototyping and machine lab. The mold and NorHA hydrogel components (NorHA, peptide crosslinker, and RGD) were sterilized by overnight lyophilization followed by germicidal UV irradiation for 2-3 hours. The LAP photoinitiator stock solution was sterile filtered prior to use. The sterilized rectangular mold was placed around the surgically created defect and a UV light (365 nm, 7 mW cm^-2^) was positioned directly above the injury. To facilitate *in situ* polymerization the UV light was turned on and the NorHA precursor solution (215 μL) was added dropwise into the injury area and irradiated for 5 mins. The mold served to retain the hydrogel precursor solution at the injury site prior to complete gelation and was removed after the first 2 mins of UV exposure. Following hydrogel delivery, the fascia was sutured back in place using 6-0 vicryl sutures and the skin was closed with 5-0 prolene using interrupted sutures. Skin glue was applied over top of the prolene sutures to avoid reopening of the injury. Following surgery, animal health was monitored daily, and buprenorphine (0.05 mg/kg) was administered subcutaneously for 3 days before sutures were removed at day 14. At 12 and 24 weeks post-surgery, the experimental and uninjured contralateral control muscles were removed for *ex vivo* functional testing and histological analysis (*n* = 8 for each experimental group).

### 2.7 Ex vivo muscle functional analysis

At terminal time points, 12 or 24 weeks after surgery, LD muscles were explanted and force generation was assessed in an *ex vivo* muscle stimulation apparatus as previously described^3,42,43^. Briefly, all explants were performed aseptically under general anesthesia. The incision site was shaved, wiped with alcohol, and a longitudinal skin incision was made over the spine of anesthetized rats. The skin and fascia were separated to expose the entirety of both LD muscles (experimental and contralateral control) for each rat from the thoracolumbar fascia to the humeral tendon. The humeral tendon was tied with 0-0 silk sutures (Ethicon) and severed from the insertion point. The proximal end was then tied, and the muscle was excised and transferred to individual chambers of a DMT 750 tissue organ bath system (DMT, Ann Arbor, MI). Each chamber was filled with Krebs–Ringer buffer solution (pH 7.4, concentrations in mM: 121.0 NaCl, 5.0 KCl, 0.5 MgCl_2_, 1.8 CaCl_2_, 24.0 NaHCO_3_, 0.4 NaH_2_PO_4_, and 5.5 glucose; Sigma) at 37°C and bubbled with 95% O_2_ and 5% CO_2_.

Excised muscles were then positioned between custom made platinum electrodes with the fascial end affixed to a stationary support and humeral tendon attached to a force transducer. Isometric muscle contraction was then quantified via direct electrical stimulation using a 0.2 ms pulse at 30 V supplied by a Grass S88 stimulator (Grass, Warwick, RI). Data visualization and acquisition were performed using Power Lab/8sp systems (AD Instruments, Colorado Springs, CO). After ensuring that muscles were securely attached and positioned between electrodes, they were subjected to a series of twitch contractions to determine optimal muscle length (*L*_0_). Force production as a function of stimulation frequency (1, 50, 100, 150, and 200 Hz) was then measured during isometric contractions (750 ms trains of 0.2 ms pulses). Following each stimulation, muscles were allowed to recover for 4 min before applying the next impulse. Force as a function of frequency curves were fit to the following dose/response equation^3,42^:

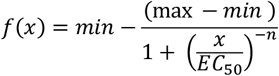

Where *x* is the stimulation frequency in Hz and *min* and *max* represent twitch force (*P*_*t*_) and maximal force of contraction (*P*_0_). *EC*_50_ is the stimulation frequency at which half of the maximum force amplitude is observed or *P*_0_ – *P*_*t*_ and *n* is the slope of the linear portion of the force/frequency curve. Force production was also normalized to animal body weight at each time point. Furthermore, specific force (specific *P*_0_) was calculated by normalizing *P*_0_ to an approximate physiological cross-sectional area (*PCSA*) using the following equation, where muscle density is 1.06 g/cm^3^:

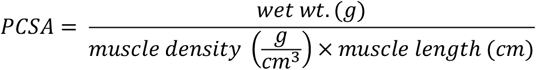

The PCSA calculation described here is an approximation for the force measurement setup previously outlined.

### 2.8 Histology analysis

Following functional testing all retrieved muscles were photographed before fixation in 4% paraformaldehyde at 4°C. Muscles were then divided into three sections: the upper third (near the humeral tendon insertion), the middle third (including some of the defect area), and the lower third (toward the thoracolumbar fascia). Upper and lower sections were used for longitudinal analysis while middle portions were used for cross-sectional analysis (Fig. S3). Muscle samples were then embedded in paraffin wax and sectioned to 5 μm thick samples. Masson’s trichrome stains were administered to 5 slides per muscle (2 sections per slide for a total of 10 individual tissue sections) using conventional techniques to analyze tissue morphology, muscle repair, and fibrotic response for at least three muscles per experimental group. Muscle sections were then visualized using a Nikon upright light microscope at various magnifications. The extent of tissue recovery following treatment was visual inspected using ImageJ across the 10 sections to gain a more complete understanding of tissue repair. The ‘zone of regeneration’ was characterized by the presence of small disorganized myofibers, a hallmark of regenerating muscle^3^. The area of regeneration was then quantified using ImageJ’s built-in measuring function at the interface of the defect and native tissue by two independent researchers (*n* = 3 muscles for each group) (Fig. S3). In the event of a greater than 50% difference between measured zones of regeneration, the analysis was repeated a third time.

### 2.9 Muscle fiber minimum Feret diameter quantification

Following histological evaluation of the muscle zone of regeneration, minimum Feret diameter, the measure of an object size along its minimum axis, and fiber cross sectional area (FCSA) were evaluated using a semi-automated image processing pipeline in ImageJ. Tissue sections corresponding to mean zone of regeneration measurements were first pre-processed using color deconvolution to separate muscle fibers. Since Masson’s trichrome stains muscle fibers in red, this channel was used for subsequent segmentation. The image was then binarized using the automated Otsu thresholding method in ImageJ and a watershed segmentation was applied to isolate individual muscle fibers. Minimum Feret diameter and FCSA were then quantified using the analyze particles tool in ImageJ. The accuracy of image processing pipeline was validated using manual counting.

### 2.10 Immunohistochemical (IHC) analysis

Unstained paraffin-embedded slides were subjected to an antigen retrieval protocol (H-3301; Vector Laboratories) to visualize vascularization and assess macrophage response to injury. To reduce autofluorescence of muscle tissue, samples were subjected to a treatment with 0.3% Sudan black solution for 10 min prior to blocking for 2 h (Dako Blocking Solution X0909; Agilent Technologies, Santa Clara, CA). Immunofluorescence staining was then performed using antibodies to detect CD31 (dilution 1:250; Novus Biologicals NB100-2284), α-smooth muscle actin (αSMA, conjugated to 488 fluorochrome, F3777, dilution 1:250; Sigma Aldrich), CD68 (1:100, MCA341R; Bio-Rad), and CD163 (1:400, ab182422; Abcam), overnight at 4°C. Next, samples were incubated for 2 hours at ambient temperature with the following secondary antibodies: Alexa Fluor 647 Fab 2 fragment goat anti-rabbit (dilution 1:500), Alexa Fluor 488 goat anti-chicken (dilution 1:600), Alexa Fluor 488 goat anti-mouse (dilution 1:400), or Alexa Fluor 594 goat anti-rabbit (dilution 1:400). Finally, slides were mounted with DAPI-containing mounting media and stored in a light protected environment at −20 °C until imaging.

All samples were imaged on a Leica DMi8 inverted confocal microscope using a 20x oil immersion objective. Immunohistochemical analyses were performed on LD muscle sections adjacent to the calculated mean zone of regeneration (*n* = 3 muscles for each group). For each sample, z-stack and tile scan capabilities were used to acquired six sequential images at the zone of regeneration defined from histological analysis. CD31 and αSMA-labeled structures were quantified for all six fields of view using the ImageJ analyzed particles tool. A custom MATLAB code was used to quantify the number of M1 macrophages stained CD68^+^/CD163^-^ and M2 macrophages stained CD68^+^/CD163^+^^44,45^.

### 2.11 Statistical Analysis

Data are presented as means and their standard deviations (SDs) unless otherwise indicated. NorHA hydrogel mechanical properties were analyzed for *n* = 3 hydrogels per group. Histological and immunohistochemical analysis was conducted for *n* = 3 muscles per group. Data normality distribution was evaluated using the D’Agostino and Pearson test. Kruskal-Wallis with Dunn’s multiple comparisons tests were performed for data that were not normally distributed (i.e., minimum Feret diameter and the zone of regeneration at 12 weeks). Functional data, rheological properties, histology, and immunohistochemical characterization were statistically analyzed by one-way analysis of variance (ANOVA). Upon finding any statistically significant differences via ANOVA, post-hoc multiple comparison tests of parameters of interest were performed using Tukey’s HSD test. Isometric force, minimum Feret diameter frequency, and FCSA frequency were statistically analyzed using paired two-way (ANOVA) with either Tukey’s or Dunnett’s multiple comparisons tests. These statistical analyses were conducted using GraphPad Prism 8.0 and R. *P* values < 0.05 were considered statistically significant.

## 3. Results

### 3.1 Norbornene-modified HA hydrogel development and characterization

Norbornene-modified hyaluronic acid (NorHA) hydrogels were formed via light-mediated thiol-ene click chemistry. This robust click chemistry allows for facile control over hydrogel mechanical properties by altering the reaction parameters including: macromer and crosslinker concentration, light intensity and duration, and degree of norbornene modification. In this study, we synthesized a matrix metalloproteinase (MMP)-degradable peptide previously shown to degrade in the presence of MMP-1 and MMP-2^41^ with pendant cysteine residues to serve as the dithiol crosslinker. NorHA hydrogel stiffness was controlled by incorporating varying amounts of peptide crosslinker to approximate the tissue mechanical properties of developing^38,39^ and adult^29^ skeletal muscle tissue. Oscillatory shear rheology confirmed that low, medium, and high concentrations of peptide crosslinker produced hydrogels with elastic moduli of 1.1 ± 0.002, 3.0 ± 0.002, and 10.6 ± 0.006 kPa, respectively (calculated as three times the storage modulus assuming a Poisson’s ratio of 0.5^20^, *n* = 3) (Fig. 1). Additionally, UV light-mediated crosslinking occurred rapidly upon irradiation (~ 20 seconds), as indicated by the dramatic increase in storage modulus (G’) and quick plateau shortly after the light was turned off. All NorHA hydrogel crosslinking was carried out using 365 nm light at an intensity of 7 mW cm^−2^ for 2 min which has previously been show to facilitate uniform crosslinking and not detrimentally affect cell viability^19,46,47^.

**Fig. 1:**
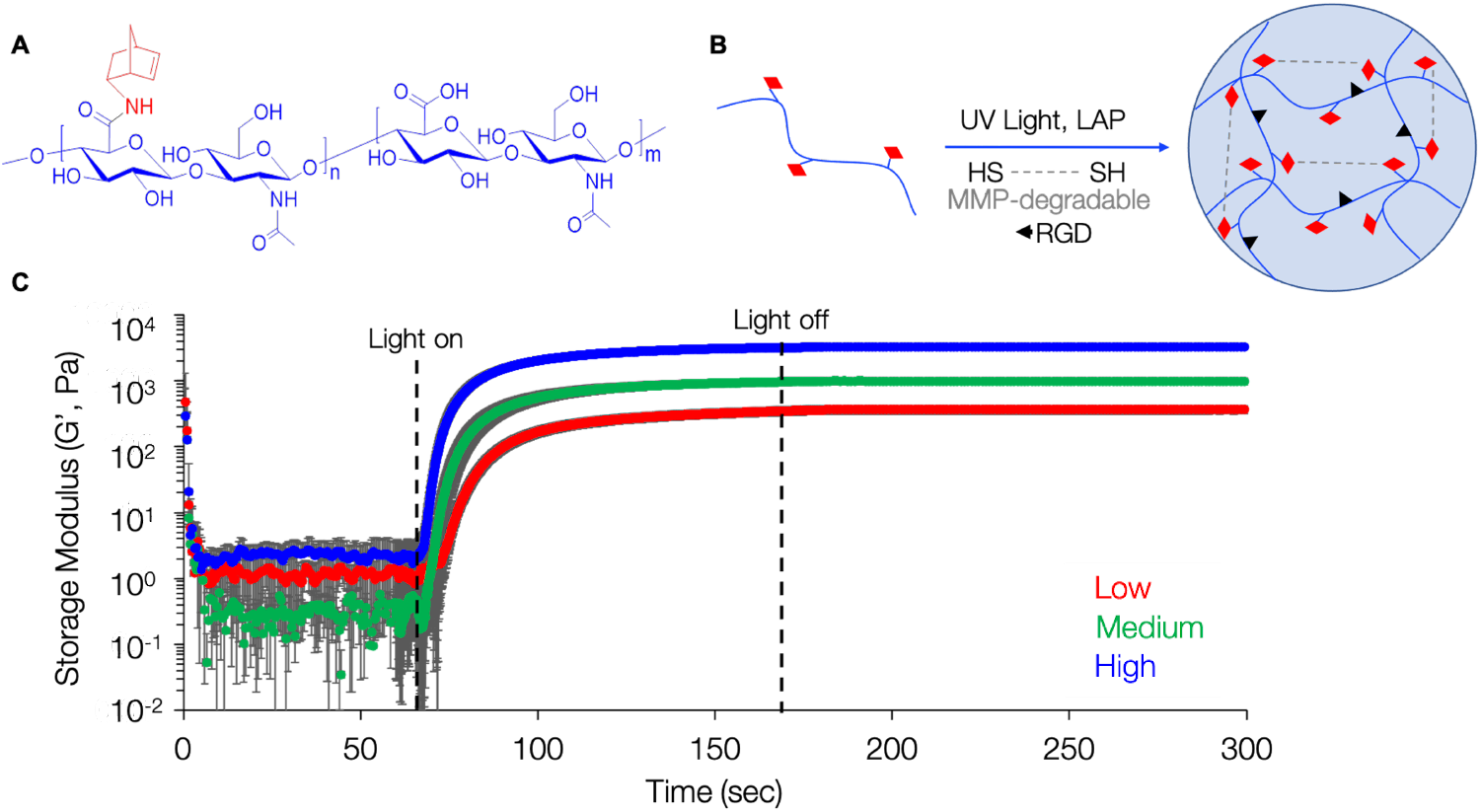
Hydrogel mechanical characterization. (A) Chemical structure of norbornene-modified hyaluronic acid (NorHA). (B) Schematic of reaction between NorHA (HA: *blue*, norbornene: *red*) and MMP-degradable dithiol peptide via UV light-mediated thiol-ene click chemistry. (C) Oscillatory shear rheology showed that increasing the concentration of MMP-degradable peptide crosslinker resulted in hydrogels of increasing stiffness. Low (1 kPa), medium (3 kPa), and high (10 kPa) elastic modulus hydrogels were developed to approximate a range of physiological muscle tissue stiffnesses. Data presented as mean +/− SD, *n* = 3 hydrogels per experimental group.

### 3.2 Creation of rat LD VML injury and hydrogel delivery

Following material characterization, NorHA hydrogels were delivered into an established rat latissimus dorsi volumetric muscle loss (VML) muscle injury model (Fig. 2A, B) via *in situ* photopolymerization. The weight and area of excised muscle were statistically similar across experimental groups, indicating creation of comparable VML injuries. Four experimental groups were investigated: No repair controls (NR, 150.5 ± 12.0 mg, 2.44 ± 0.22 cm^2^) and hydrogel groups with low (146.4 ± 10.8 mg, 2.27 ± 0.19 cm^2^), medium (148.0 ± 8.51 mg, 2.35 ± 0.27 cm^2^), and high (148.6 ± 10.7 mg, 2.48 ± 0.20 cm^2^) stiffness (**Fig. 2C, D**). The final defect area was slightly larger than the initial measured region (1.5 × 1.2 cm) because after the tissue was excised the remaining muscle fibers continued to be under tension, resulting in strain at the injury site and ultimately a larger defect. Following creation of the injury, NorHA hydrogel solutions were photopolymerized within the defect area and secured in place by suturing the fascia and skin.

**Fig. 2:**
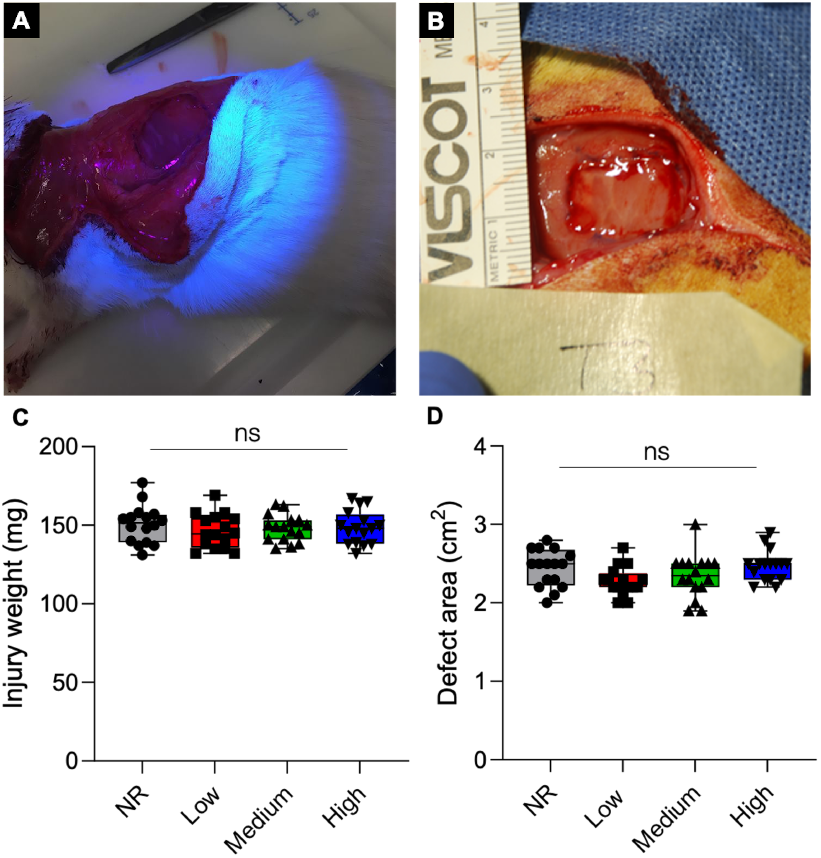
*In situ* photopolymerization enables facile and reproducible hydrogel delivery to rat LD VML defects. (A) NorHA hydrogels were photopolymerized *in situ* and (B) conformed to the injury dimensions, completely filling the defect area. (C) Surgical dissections resulted in reproducible injuries of 148.4 ± 10.4 mg and (D) 2.44 ± 0.22 cm^2^ that were not statistically different between experimental groups. Data presented as box plots of interquartile range (line: median) with whiskers showing minimum and maximum values, *n* = 16 animals per experimental group.

### 3.3 Ex vivo functional assessment of LD force generation

Functional muscle recovery was assessed at 12 and 24 weeks post-surgery via *ex vivo* electrical stimulation on experimental and contralateral control muscles. Briefly, LD muscles were excised and attached to a force transducer in a recirculating organ bath. Isometric contractile force in response to direct muscle electrical stimulation was recorded over a range of frequencies (1-200 Hz). After 12 weeks, the mean maximal isometric contraction following surgical creation of VML injuries was significantly reduced compared to contralateral control muscles across all experimental groups (Fig. 3B). In non-repaired VML injuries, maximal contraction was reduced to 64 ± 18% of contralateral control tetanic contraction force (Fig. S4). Similarly, stimulation of muscles in which high stiffness hydrogels (10 kPa) were implanted at the site of VML injury, mimicking the stiffness of adult skeletal muscle tissue, resulted in 64 ± 19% of the mean maximal isometric force production of the uninjured contralateral control muscle. While not statistically greater than the no repair (NR) group, medium stiffness hydrogels (3 kPa), approximating a more compliant microenvironment, resulted in the greatest force recovery at 12 weeks producing 82 ± 13% of contralateral control values. Furthermore, when maximal isometric contraction was normalized to animal body weight at the time of explant, maximal force generation of medium stiffness hydrogels (6.23 ± 1.01 mN/g, *n* = 8) was statistically similar to native contralateral responses (7.61 ± 1.20 mN/g, *n* = 32) (Fig. S4C). In contrast, all other experimental groups maintained a significant reduction in force generation compared to the native LD when normalized to animal body weight; although the specific force (i.e., specific P_0_) was similar to contralateral control values across all treatment groups except the group implanted with the high stiffness (10 kPa) hydrogel (Fig. 3C). Not surprisingly, evaluation of explanted LD muscle mass and size revealed that NR muscles were significantly lighter and had reduced approximate physiological cross-sectional area (PCSA) calculated from measured muscle weight, length, and density (**Table 1)**when compared to hydrogel treated and contralateral muscles (**Fig. 3D**). Furthermore, hydrogel treated muscles had statistically similar mass and PCSA to contralateral control muscles. Following *ex vivo* functional testing, images of retrieved LD muscles were taken for each treatment group (**Fig. S5**). Qualitative analysis of explanted LD muscles indicated that NorHA hydrogels were enzymatically degraded in low stiffness treated animals within 12 weeks although some hydrogel remained in medium and high stiffness groups. Additionally, the hydrogel groups appeared to integrate with native tissue while the NR group appeared to lack new tissue formation and instead contained a thin layer of fibrous tissue.

**Fig. 3:**
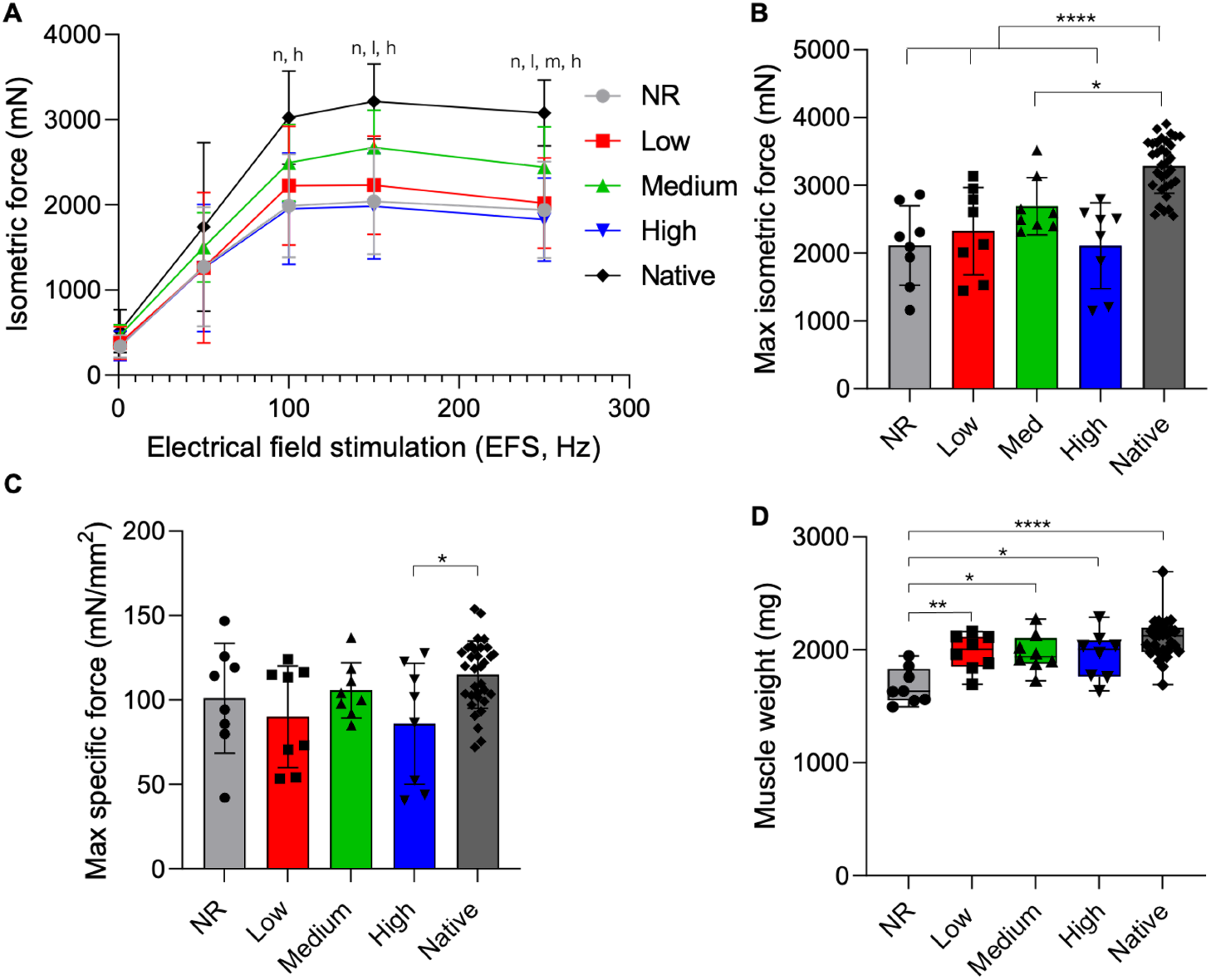
Medium stiffness hydrogels support muscle force generation most similar to native tissue. (A) Dose/response curve and (B) maximum isometric contraction force in response to electrical field stimulation at 12 weeks post-surgery. Statistically significant differences from native (*P* < 0.05) are denoted by *n* (NR), *l* (low), *m* (medium), and *h* (high). (C) Specific force (specific P_0_), calculated by normalizing maximal isometric force (P0) to physiological cross-sectional area (PCSA), was similar to contralateral control values across all treatment groups except muscles treated with the high stiffness (10 kPa) hydrogels. (D) Muscle weight at the time of explant shows no repair (NR) muscles were significantly lighter than other experimental groups. Data presented as mean +/− SD (panels A-C) while panel D data are presented as box plots of interquartile range (line: median) with whiskers showing minimum and maximum values. * *P* < 0.05, ** *P* < 0.01, *** *P* < 0.001, **** *P* < 0.0001. *n* = 8 muscles per experimental group and *n* = 32 for the native contralateral control.

**Table 1:**
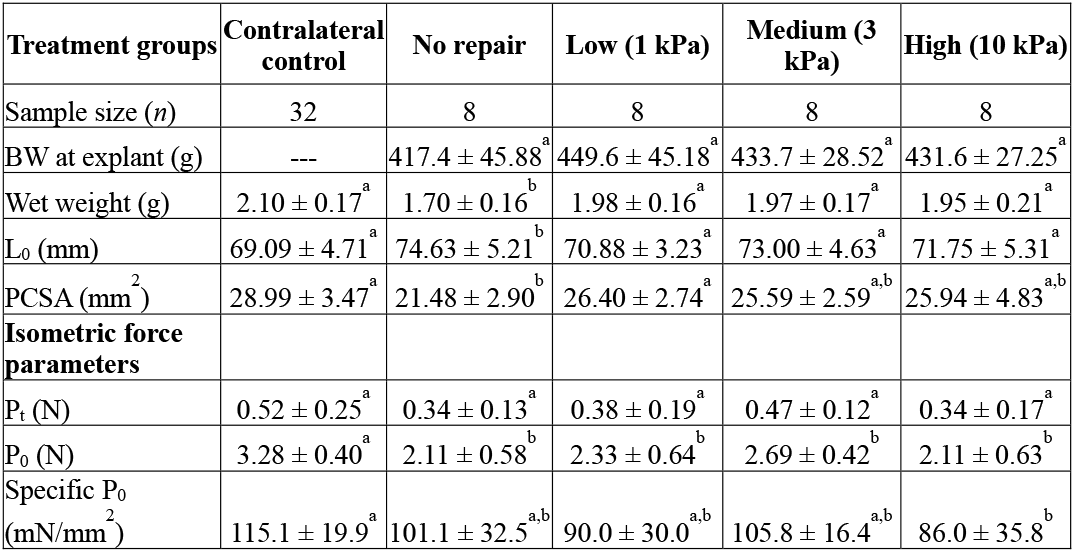
Summary of rat latissimus dorsi muscle functional measurement data at 12 weeks post-injury

Values are presented as mean ± standard deviation. Statistical differences were analyzed using Tukey’s post-hoc test after performing a one-way ANOVA. Values with the same letter were not significantly different (*P* > 0.05), while values with dissimilar letters were significantly different (*P* < 0.05). Peak twitch response (Pt) and maximal isometric contraction force (P0) were measured by *ex vivo* direct electric stimuli (0.2 ms, 30 V). Measured P_0_ was normalized to PCSA to determine specific force (specific P_0_). BW, body weight; ---, analysis not applicable to contralateral control; L0, muscle length; PCSA, physiological cross-sectional area.

To assess longer term functional recovery *ex vivo* electrical stimulation of explanted muscles was also assessed at 24 weeks post-surgery. Mean maximal isometric contraction of NR muscles remained significantly reduced at the later time point and resulted in 73 ± 17% of contralateral control force production (**Fig. 4, Fig. S4**). NR muscles also produced significantly less isometric force across various stimulation frequencies when compared to contralateral control muscles. Similarly, the addition of high stiffness hydrogels to injured muscles resulted in minimal functional recovery of muscle force that was significantly reduced when compared to the contralateral controls (77 ± 18%). Low and medium stiffness hydrogel treated muscles resulted in maximal tetanic forces that were statistically similar to contralateral control muscles and recovered to 83 ± 14% and 90 ± 18% of native values respectively. The same trends were observed when maximal isometric contraction was normalized to animal body weight (**Fig. S4**). While lower stiffness hydrogels supported improved functional recovery, PCSA was statistically reduced across all experimental groups and muscle weight was reduced in low and NR muscles when compared to contralateral control tissue (**Table 2**). Similar to results observed at 12 weeks, evaluation of the specific force (i.e., specific P_0_) showed similar levels of force generation across all treatment groups when compared to contralateral control values. Also consistent with data at 12 weeks, qualitative analysis of gross tissue morphology at 24 weeks showed that NR muscle injuries were covered in a thin layer of connective tissue (**Fig. S5**). While hydrogel groups largely appeared to integrate with host tissue, there were remnants of NorHA hydrogel observed in some medium and high stiffness hydrogel treated muscles.

**Fig. 4:**
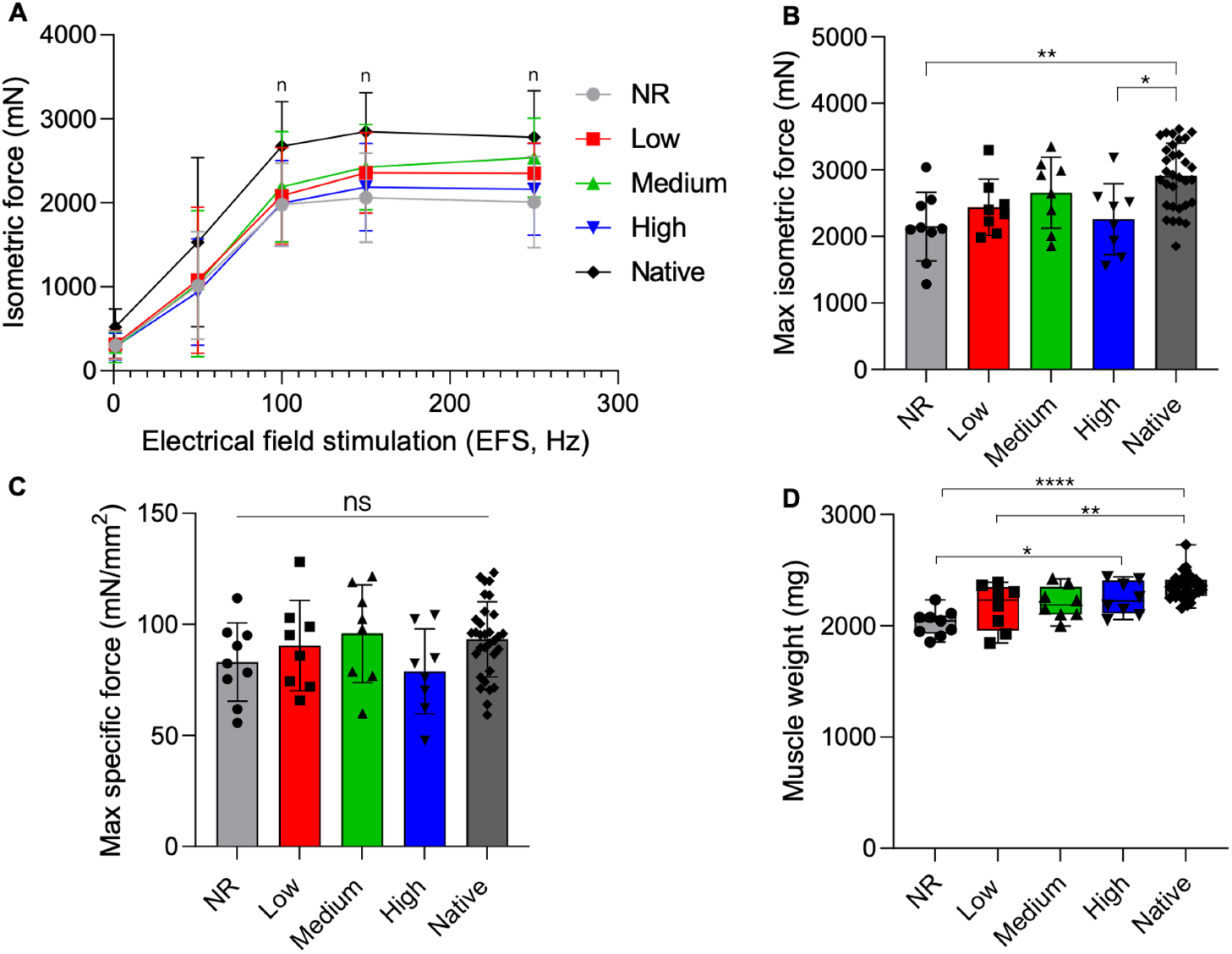
Lower stiffness hydrogels enable similar force generation to native muscle at 24 weeks. (A) Dose/response curve and (B) maximum isometric contraction force in response to electrical field stimulation at 24 weeks post-surgery. Statistically significant differences from native (*P* < 0.05) are denoted by *n* (NR). (C) Specific P_0_ was similar to contralateral control values across all treatment groups at the time of explant. (D) Explanted muscle weight shows no repair (NR) and low stiffness hydrogel treated muscles were significantly lighter than native muscle. Data presented as mean +/− SD (panels A-C) while panel D data are presented as box plots of interquartile range (line: median) with whiskers showing minimum and maximum values. * *P* < 0.05, ** *P* < 0.01, *** *P* < 0.001, **** *P* < 0.0001. *n* = 8 muscles per experimental group and *n* = 33 for the native contralateral control.

**Table 2:** Summary of rat latissimus dorsi muscle functional measurement data at 24 weeks post-injury

| Treatment groups | Contralateral control | No repair | Low (1 kPa) | Medium (3 kPa) | High (10 kPa) |
| --- | --- | --- | --- | --- | --- |
| Sample size (n) | 33 | 9 | 8 | 8 | 8 |
| BW at explant (g) | -- | 491.3 ± 35.53<sup>a</sup> | 485.9 ± 22.58<sup>a</sup> | 499.8 ± 36.71<sup>a</sup> | 483.0 ± 27.89<sup>a</sup> |
| Wet weight (g) | 2.10 ± 0.17<sup>a</sup> | 2.02 ± 0.12<sup>b</sup> | 2.17 ± 0.21<sup>b,c</sup> | 2.21 ± 0.14<sup>a,b</sup> | 2.25 ± 0.15<sup>a,c</sup> |
| L<sub>0</sub> (mm) | 71.13 ± 4.05<sup>a</sup> | 74.22 ± 3.46<sup>a</sup> | 74.88 ± 3.40<sup>a</sup> | 74.50 ± 3.46<sup>a</sup> | 73.75 ± 4.89<sup>a</sup> |
| PCSA (mm<sup>2</sup>) | 31.30 ± 1.80<sup>a</sup> | 25.78 ± 1.74<sup>b</sup> | 27.40 ± 3.25<sup>b,c</sup> | 27.96 ± 1.97<sup>b,c</sup> | 28.86 ± 2.48<sup>c</sup> |
| **Isometric force parameters** |  |  |  |  |  |
| P<sub>t</sub> (N) | 0.52 ± 0.21<sup>a</sup> | 0.30 ± 0.17<sup>b</sup> | 0.32 ± 0.17<sup>a,b</sup> | 0.28 ± 0.18<sup>b</sup> | 0.29 ± 0.16<sup>b</sup> |
| P<sub>0</sub> (N) | 2.94 ± 0.46<sup>a</sup> | 2.15 ± 0.51<sup>b</sup> | 2.44 ± 0.42<sup>a,b</sup> | 2.66 ± 0.53<sup>a,b</sup> | 2.26 ± 0.53<sup>b</sup> |
| Specific P<sub>0</sub> (mN/mm<sup>2</sup>) | 94.4 ± 15.9<sup>a</sup> | 83.1 ± 17.6<sup>a</sup> | 90.5 ± 20.3<sup>a</sup> | 95.9 ± 22.0<sup>a</sup> | 78.9 ± 19.1<sup>a</sup> |

Values are presented as mean ± standard deviation. Statistical differences were analyzed using Tukey’s post-hoc test after performing a one-way ANOVA. Values with the same letter were not significantly different (*P* > 0.05), while values with dissimilar letters were significantly different (*P* < 0.05). Peak twitch response (Pt) and maximal isometric contraction force (P0) were measured by *ex vivo* direct electric stimuli (0.2 ms, 30 V). Measured P_0_ was normalized to PCSA to determine specific force (specific P_0_). BW, body weight; ---, analysis not applicable to contralateral control; L0, muscle length; PCSA, physiological cross-sectional area.

### 3.4 Histological analysis

After functional testing, explanted muscles were formalin fixed and paraffin embedded for histological analysis. Three representative samples from each treatment group were transversely sectioned and underwent Masson’s trichrome staining to assess muscle fiber morphology and fibrotic response to VML injury. Stained tissue samples were analyzed to identify the interface area between the injury and the remaining native muscle tissue. The ‘zone of regeneration’, characterized by the presence of smaller more disorganized muscle fibers, was used to quantify the extent of tissue recovery across experimental groups as previously described3,48. As observed from the gross tissue morphology, the VML defect area was easily identified across experimental groups. The injured tissues exhibited a clear lack of muscle volume reconstitution and instead consisted of a thin membrane of connective tissue that filled the surgical defect. Quantitative analysis of LD muscle sections showed statistically similar zones of regeneration across all experimental groups at 12 weeks post-surgery (Fig. S6, data reported as the median, 1^st^, and 3^rd^ quartiles; NR: 0.18, 0.16, 0.22 mm2; Low: 0.38, 0.24, 0.55 mm^2^; Medium: 0.32, 0.30, 0.50 mm^2^; High: 0.25, 0.21, 0.35 mm^2^; *P* > 0.05). However, hydrogel treated muscles showed some evidence of muscle recovery as indicated by disorganized myofiber infiltration.

To assess if NorHA hydrogels supported longer term functional recovery, the same zone of regeneration quantification scheme was repeated for injured muscles at 24 weeks post-surgery (**Fig. 5**). Similarly, NR treated muscles displayed a thin sheet of connective tissue and a clear absence of regenerating muscle fibers (NR zone of regeneration: 0.16 ± 0.02 mm^2^) characteristic of a VML injury. High stiffness hydrogel treated muscles also resulted in minimal tissue recovery and was characterized by the presence of fibrotic tissue (High: 0.13 ± 0.03 mm^2^). Low stiffness hydrogel treated tissues showed a modest increase in the zone of regeneration when compared to NR muscles (low: 0.30 ± 0.08 mm^2^), although these results were not statistically significant (*P* = 0.09). However, the zone of regeneration in low stiffness hydrogel treated muscles was significantly larger than high stiffness hydrogel treated tissues. In contrast, medium stiffness hydrogel treated muscles showed reduced levels of fibrosis and improved myogenesis as indicated by myofiber infiltration into the wound bed, reduced connective tissue, and a significantly larger zone of regeneration (medium: 0.44 ± 0.08 mm^2^) when compared to NR and high stiffness hydrogel treated tissues. Ultimately, these results corroborate the isometric force data indicating that muscle regeneration is critical for functional muscle recovery.

**Fig. 5:**
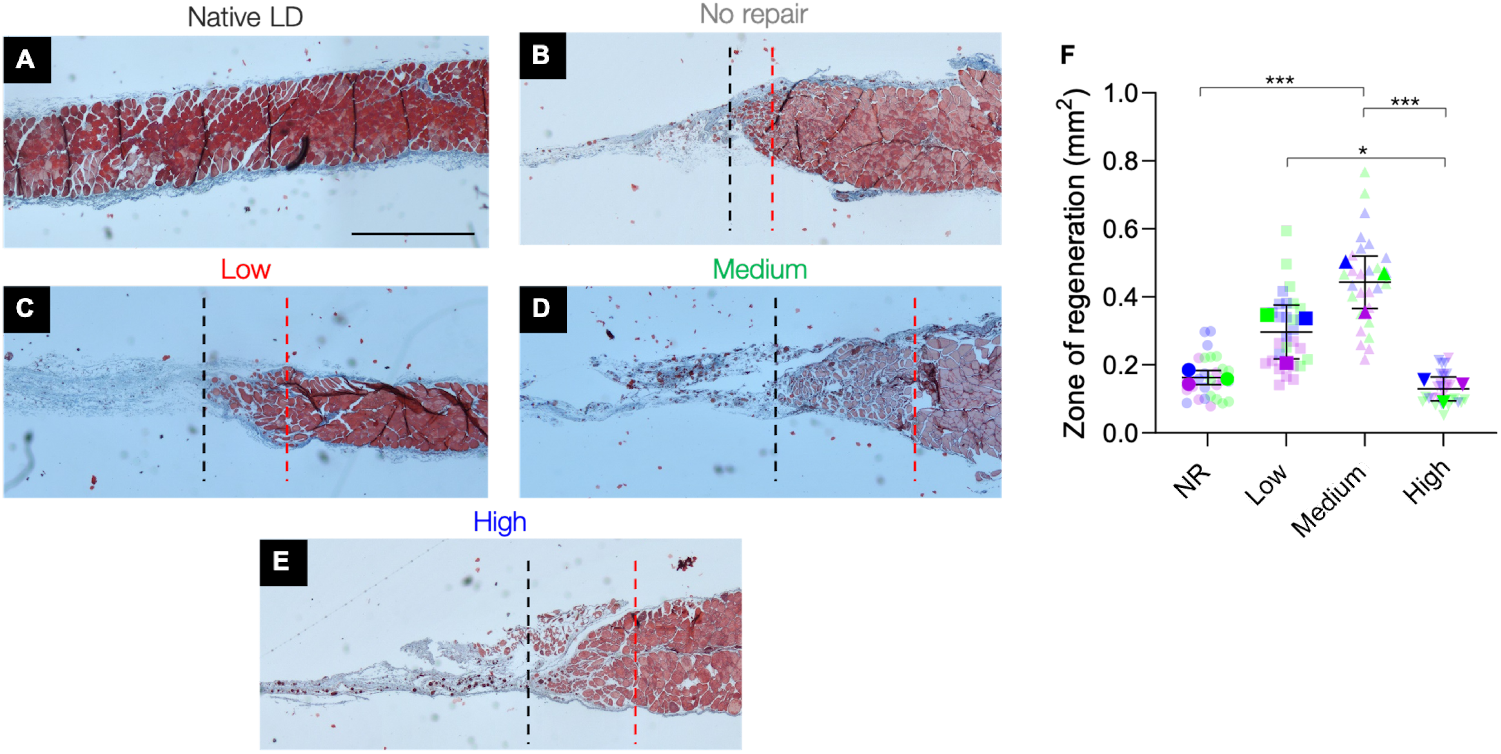
Histological analysis of LD muscles shows improved myogenesis in medium stiffness hydrogel treated muscles at 24 weeks. (A-E) Tissue and cell morphology at the interface between native muscle and the VML injury was visualized by Masson’s trichrome staining (muscle tissue in red, collagen deposition in blue, and nuclei in black). The area between the dashed red and black lines represents the zone of regeneration characterized by small disorganized myofibers. The red line denotes the interface between native muscle and VML injury site and the black line represents the last point at which detectable myofibers were present. (F) Quantification of the zone of regeneration at 24 weeks post-injury indicated improved myogenesis in medium stiffness hydrogel treated muscles. Colors denote different experimental muscles (biological replicates, *n* = 3 per group) with solid shapes representing the mean and translucent shapes representing individual sections per muscle (technical replicates, *n* = 10). Data presented as mean +/− SD. * *P* < 0.05, ** *P* < 0.01, *** *P* < 0.001. Scale bar: 1 mm.

### 3.5 Characterization of regenerating muscle fibers

Following analysis of the zone of regeneration, regenerating muscle fibers at the implanted hydrogel-native tissue interface were further characterized. Specifically, the number of muscle fibers as well as the fiber cross-sectional area (FCSA, Fig. S7) and minimum Feret diameter (the measure of an object size along its minimum axis), were quantified in the zone of regeneration for each experimental group (Fig. 6, Fig. S8). These findings were then compared to muscle fiber characteristics in the native contralateral control muscles. Medium stiffness hydrogel treated tissues contained significantly more muscle fibers in the zone of regeneration (189.7 ± 54.5 per section imaged) when compared to NR muscles (88.3 ± 15.6). Similarly, low (130.7 ± 7.2) and high stiffness hydrogel treated muscles (128.0 ± 25.5) trended toward larger quantities of muscle fibers when compared to NR tissues, but these results were not statistically significantly different. Further, analysis of FCSA and minimum Feret diameter revealed that regenerating fibers in the zone of regeneration were smaller than muscle fibers in native uninjured tissue regardless of treatment type. This trend is apparent by the increased frequency of smaller diameter fibers and decreased presence of larger muscle fibers when compared to fibers in native tissues. In particular, NR and high stiffness hydrogel treated muscles displayed a significantly reduced frequency of medium sized muscle fibers when compared to uninjured tissues. Quantification of the median minimum Feret diameter showed that fiber size was significantly reduced in NR (21.5, 16.2, 28.3 μm; reported as median, 1^st^, and 3^rd^ quartiles) treated tissues when compared to native muscle (38.4, 28.0, 51.3 μm). In contrast, low (28.4, 19.7, 42.2 μm), medium (27.9, 20.1, 37.7 μm), and high (23.1, 16.6, 31.0 μm) stiffness hydrogel treated muscles contained statistically similar median fiber diameters compared to uninjured muscle; however, overall fiber size was reduced.

**Fig. 6:**
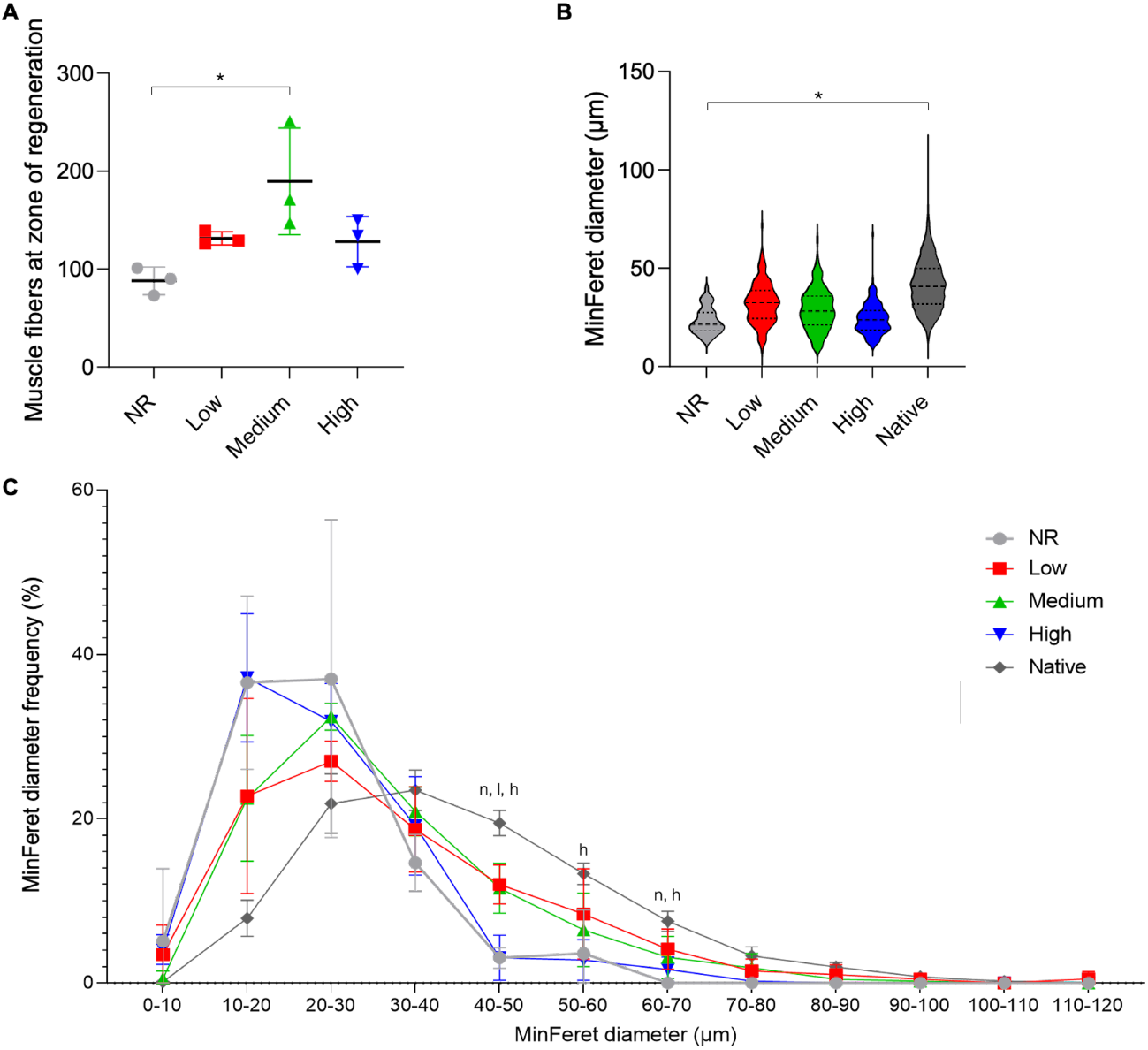
Medium stiffness hydrogels support increased number of muscle fibers. (A) Quantification of the number of muscle fibers in the zone of regeneration per muscle section revealed that medium stiffness hydrogels supported increased myogenesis. (B) Median minimum Feret diameter (dashed lines) was significantly reduced in no repair (NR) treated muscle compared to native muscle (dotted lines, 1^st^ and 3^rd^ quartiles). (C) Minimum Feret diameter frequency distribution curve shows a leftward shift toward smaller muscle fibers compared to uninjured muscle regardless of treatment type. Statistically significant differences from native (*P* < 0.05) are denoted by *n* (NR), *l* (low), and *h* (high). Data presented as mean +/− SD (panels A, C). * *P* < 0.05. *n* = 3 muscles per experimental group.

### 3.6 Assessment of vascularization

Following histological analysis, immunohistochemical (IHC) staining was used to probe for vascularization of 24-week muscle samples in the zone of regeneration. The presence of endothelial cells was evaluated by CD31 staining, while larger vessels were detected by the co-expression of CD31 and alpha smooth muscle actin (αSMA), a marker of pericytes. The presence of αSMA was significantly reduced across all experimental groups when compared to native muscle tissue, indicating reduced levels of mature vasculature. However, lower stiffness hydrogel treated muscles (low and medium) appeared to support modest levels of vascularization as indicated by the presence of CD31+ cells that was not statistically different from native muscle (low: *P* = 0.90, medium: *P* = 0.20) (Fig. 7). Similarly, retrieved muscle tissues from the NR group contained statistically similar levels of CD31+ cells but were somewhat reduced when compared to native muscle (NR: *P* = 0.057). High stiffness hydrogel treated muscles possessed significantly reduced CD31^+^ cells when compared to native tissue (high: *P* = 0.049).

**Fig. 7:**
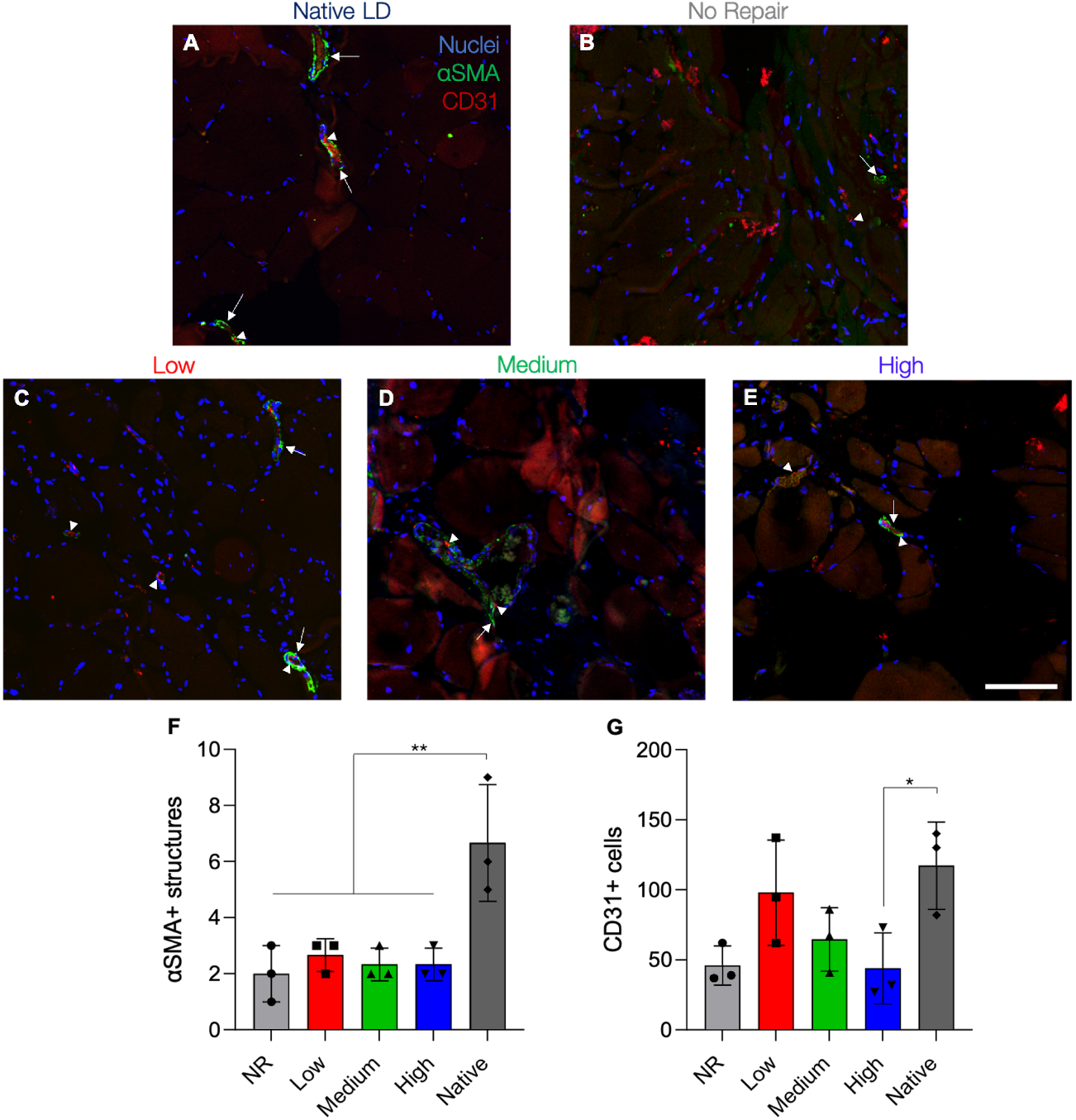
Lower stiffness hydrogels support modest vascularization 24 weeks post-VML. (A-E) Representative images of vascular staining at 24 weeks post-VML. Images were taken in the zone of regeneration showing αSMA structures (*green*, arrow) and CD31 cells (*red*, arrowhead) at the defect-tissue interface. (F) Quantification of αSMA structures indicative of arteries and veins was significantly reduced across all experimental groups compared to native muscle. (G) Lower stiffness hydrogels (1 and 3 kPa) supported statistically similar levels of CD31^+^ cells to native muscle. Data presented as mean +/− SD. * *P* < 0.05, ** *P* < 0.01. *n* = 3 muscles per experimental group. Scale bar: 100 μm.

### 3.7 Evaluation of long-term macrophage polarization

VML injuries are characterized by a prolonged inflammatory response that triggers excessive fibroblast activation, collagen deposition, ECM stiffening, and ultimately poor regenerative outcomes^31^. To assess this prolonged inflammatory cascade macrophage infiltration and polarization were evaluated in muscle sections 24 weeks post-VML (Fig. 8). The total macrophage presence in the zone of regeneration was characterized using CD68, a pan macrophage marker, and CD163, a marker of M2 macrophages, staining of three LD muscles per experimental group45. Co-expression of CD68 and CD163 indicative of alternatively activated M2 macrophages (CD68^+^/CD163^+^) was not statistically different across experimental groups^49–51^. However, the number of CD68^+^/CD163^−^ macrophages, indicative of a classically activated M1 pro-inflammatory response, remained elevated in no repair and high stiffness hydrogel treated muscles when compared to contralateral control muscles, signifying a chronic inflammatory response^52,53^.

**Fig. 8:**
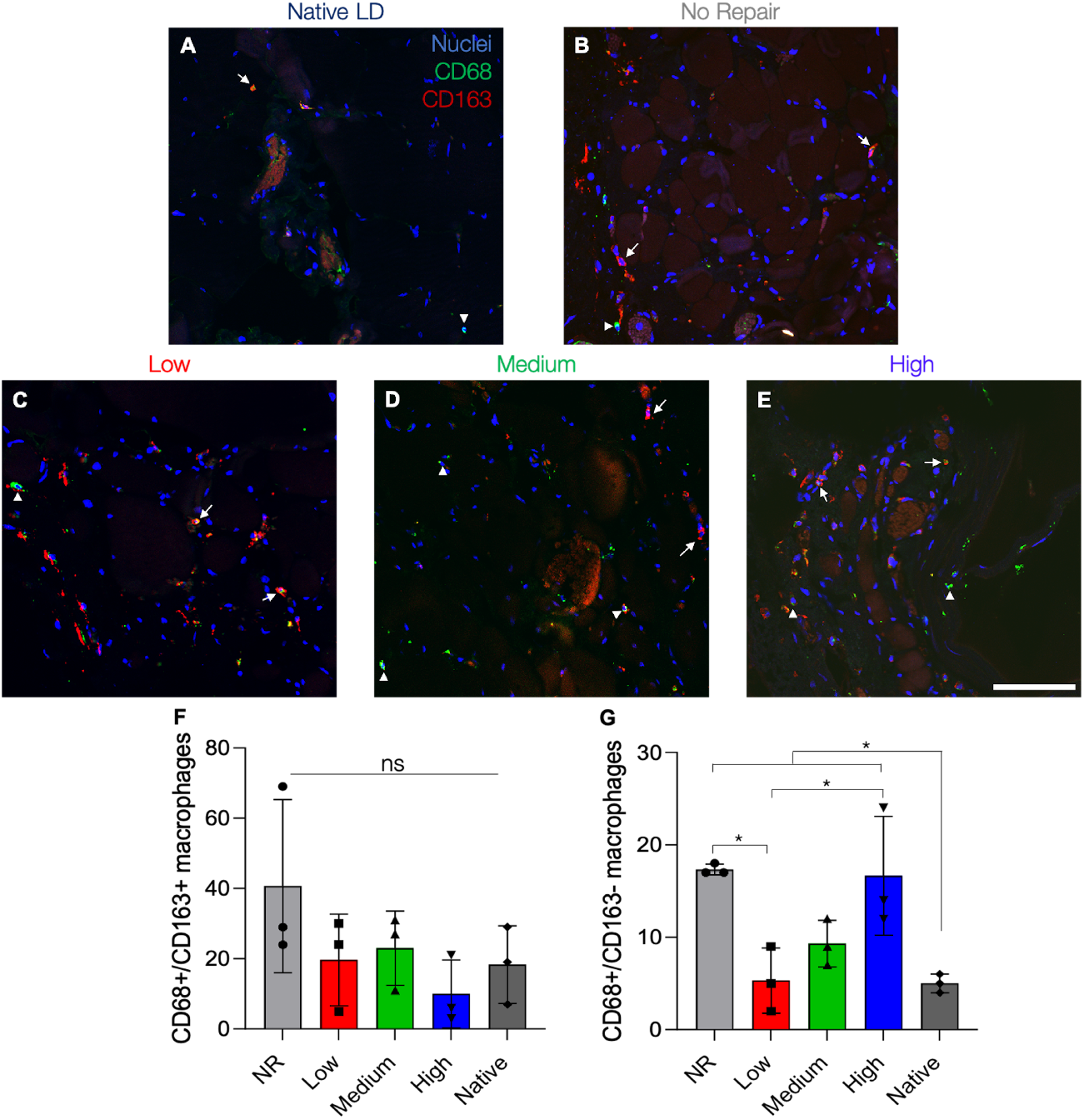
Lower stiffness hydrogels support similar macrophage polarization profiles to native muscle. (A-E) Representative images of macrophage infiltration at 24 weeks post-VML. Images were taken in the zone of regeneration showing CD68 (*green*, arrowhead) and CD163 (*red*, arrow) at the defect-tissue interface. (F) Quantification of CD68^+^/CD163^+^ cells indicative of M2 macrophages showed no statistically significant differences across experimental groups. (G) CD68^+^/CD163^−^ M1 macrophages remained significantly elevated in NR and high stiffness treated muscle indicating a prolonged inflammatory response. Data presented as mean +/− SD. * *P* < 0.05. *n* = 3 muscles per experimental group. Scale bar: 100 μm.

## 4. Discussion

Maximal functional recovery of VML injuries requires restoration of muscle volume without fibrosis and recovery of tissue function approaching preinjury levels. Despite recent advancements there remains significant challenges toward the development of more effective tissue engineering technologies for muscle repair. As a result, to begin to better inform the design criteria for improved biomaterial scaffolds for skeletal muscle tissue engineering, it is critical to understand which biophysical properties are important drivers of cell-mediated repair and which signals propagate aberrant wound healing.

While stem cell culture on 2D substrates has dramatically improved our understanding of how mechanical signals drive myogenesis^27,30^, it remains unclear if these findings translate to successful wound healing *in vivo*. To address this limitation we explored how hydrogel stiffness, a key biomaterial property, impacted skeletal muscle wound healing in a biologically relevant VML injury model.

We chose hyaluronic acid (HA) for our hydrogel platform due to its innate biological activity as a component of the ECM as well as its biocompatibility, amenability to chemical modification, and its role in promoting angiogenesis, muscle repair, and tissue development^34,54–56^. Furthermore, previous work by our lab found that degradable HA-based hydrogels could support regeneration of muscle volume, functional recovery, and improved vascularization in a rat tibialis anterior VML injury model^15^. While clearly illustrating the benefits of HA hydrogels for tissue engineering, this study did not explore how material biophysical properties including substrate elasticity impact skeletal muscle repair. As a logical extension of this prior work, the present study explored utilization of a distinct HA hydrogel that was functionalized with norbornene groups that allowed for precise control over material biophysical properties via light-mediated thiol-ene click chemistry. This chemistry was advantageous for hydrogel delivery because click reactions are characterized by their high thermodynamic driving forces and yields, generation of minimal side products, and reaction specificity^57^. As a result, polymerization of NorHA hydrogels occurred rapidly over approximately 20 seconds (**Fig. 1**), enabling localization of the hydrogel to the VML wound site without leaking. This is in stark contrast to other hydrogel crosslinking reactions which can occur over significantly longer time scales^58,59^. Additionally, *in situ* photopolymerization also allowed the hydrogel to conform to the dimensions of the injury, ensuring complete coverage of the defect (**Fig. 2**). These findings suggest that this hydrogel system can be applied to treat more geometrically complex and irregular injuries.

Skeletal muscle is a highly dynamic tissue, often responding to external stimuli to adapt, repair, and develop and become stronger^1^. However, in the context of VML and aging the skeletal muscle microenvironment stiffens, resulting in poor regenerative outcomes and loss of muscle strength due to excessive collagen deposition and fibrosis^31,40^. To support improved regenerative outcomes, we hypothesized that a microenvironment recapitulating developmental muscle mechanics would provide appropriate stimuli for muscle repair post-VML. A substrate elasticity of 10-12 kPa (matching adult muscle stiffness^29^) has previously been shown to facilitate myogenic cell differentiation^27,29^ and improved cell engraftment in an adult murine model^30^. In contrast, while the elasticity of developmental skeletal muscle tissues remains ill defined, previous studies reported stiffnesses on the order of 0.5-2 kPa^38,39^. As a result, we designed hydrogels with an elastic modulus of 10 kPa to represent healthy adult skeletal muscle stiffness, and hydrogels of 1 and 3 kPa to approximate developmental skeletal muscle microenvironments. While these hydrogel mechanical properties may mimic developmental muscle, further analysis of embryonic tissue biophysical properties would greatly inform subsequent biomaterial designs and possibly lead to superior regenerative outcomes.

After defining the mechanical targets for our hydrogel system, we chose to synthesize GCNSVPMSMRGGNCG as our peptide crosslinker owing to its previous use in proteolytically dynamic hydrogel systems^41,47^ and available pendant cysteine residues allowing for thiol-ene click gelation^19^. Previous work using poly(ethylene glycol) (PEG) showed that hydrogels crosslinked with this sequence degraded rapidly upon exposure to MMP-1 and MMP-2 as compared to other protease-degradable peptides^41^. This sequence has also been used to modulate substrate stiffness in NorHA hydrogels and to probe mechanosensitive cell signaling^47^. NorHA hydrogels formed with this peptide crosslinker were previously shown to degrade in the presence of both MMP-2 and collagenase^47^. Human mesenchymal stromal cells (hMSCs) encapsulated within MMP-degradable NorHA hydrogels were able to enzymatically remodel their local microenvironment and displayed increased spreading and nuclear localization of mechanosensitive transcriptional co-activators compared to cells within non-degradable controls^47^. In this study we were able to synthesize hydrogels with a range of material elasticities by incorporating varying amounts of the degradable peptide crosslinker.

After optimizing our material system, we aimed to explore how mechanically tunable NorHA hydrogels affected wound healing in a biologically relevant VML model. A number of different *in vivo* VML injury models have been explored in the literature with the majority of studies focusing on repair of hindlimb muscles in rodents^42,44,60,61^. These models provide a reasonable analog to combat-related injuries and extremity trauma but relatively few studies have explored muscle defects in craniofacial tissues. One such muscle, the orbicularis oris muscle (OOM), is damaged in clefts of the lip (CL) or palate and results in a VML-like injury that often requires multiple repair surgeries^62^. CL is also one of most common congenital craniofacial defects affecting 1 in 1,600 children in the United States^63^. However, many VML models fail to explore wound healing in comparatively thin, sheet-like tissues that are characteristic of craniofacial muscles such as OOM. Additionally, craniofacial tissues are markedly different from extremity muscles in both fiber composition and architecture, making comparisons with previous work challenging^64,65^. In an effort to address this knowledge gap we used a rat LD VML injury model that is dimensionally similar to the OOM muscle and contains comparable fiber type, tissue thickness, and fiber orientation^3,66^. In this biologically relevant VML injury model, our results showed that intermediate stiffness hydrogels with elastic moduli of 3 kPa facilitated improved functional muscle recovery as measured by force generation compared to non-treated muscle (NR), low, and high stiffness hydrogel variants (1 and 10 kPa respectively) (**Fig. 3**). Moreover, there was evidence for persistent muscle function up to 24 weeks post-injury, indicating that recovery may be stable over time (**Fig. 4**).

Interestingly our results showed that high stiffness hydrogels (10 kPa), which mimic adult skeletal muscle stiffness and have previously been show to support myogenic differentiation and engraftment^27,29,30^, resulted in the lowest amount of functional recovery and tissue repair of the three hydrogel groups. We believe one reason for this result is the difference in culture dimensionality as much of the previous work in the field explored cellular mechanotransduction on 2D substrates that do not accurately recreate the complexity of the 3D microenviorment^27,30^. However, a potential limitation of our current material system is that hydrogel stiffness is directly related to the number of crosslinks within the material. As a result, the increased crosslinking density required to make stiffer hydrogels may have limited cell infiltration and remodeling. While it is difficult to decouple hydrogel mechanics from degradation kinetics, future work could investigate hydrogels with constant crosslinking density and variable macromer concentration or microporous hydrogels to improve cell infiltration post-VML. In contrast, muscles treated with low stiffness hydrogels (1 kPa) facilitated improved functional recovery when compared to NR and high stiffness groups but did not reach the levels of force generation measured in medium stiffness hydrogel (3 kPa) treated muscles or contralateral control muscles (**Fig. 4**), although these results were not statistically significant. These disparate outcomes are likely due to more rapid hydrogel degradation resulting in enhanced cell infiltration that was not observed in high stiffness groups. However, the improved results observed in medium stiffness hydrogel treated muscles suggest that there may be an optimal biomaterial stiffness to facilitate adequate cell infiltration and remodeling while also modulating cell differentiation and functional repair post-VML (**Table 3**).

**Table 3:**
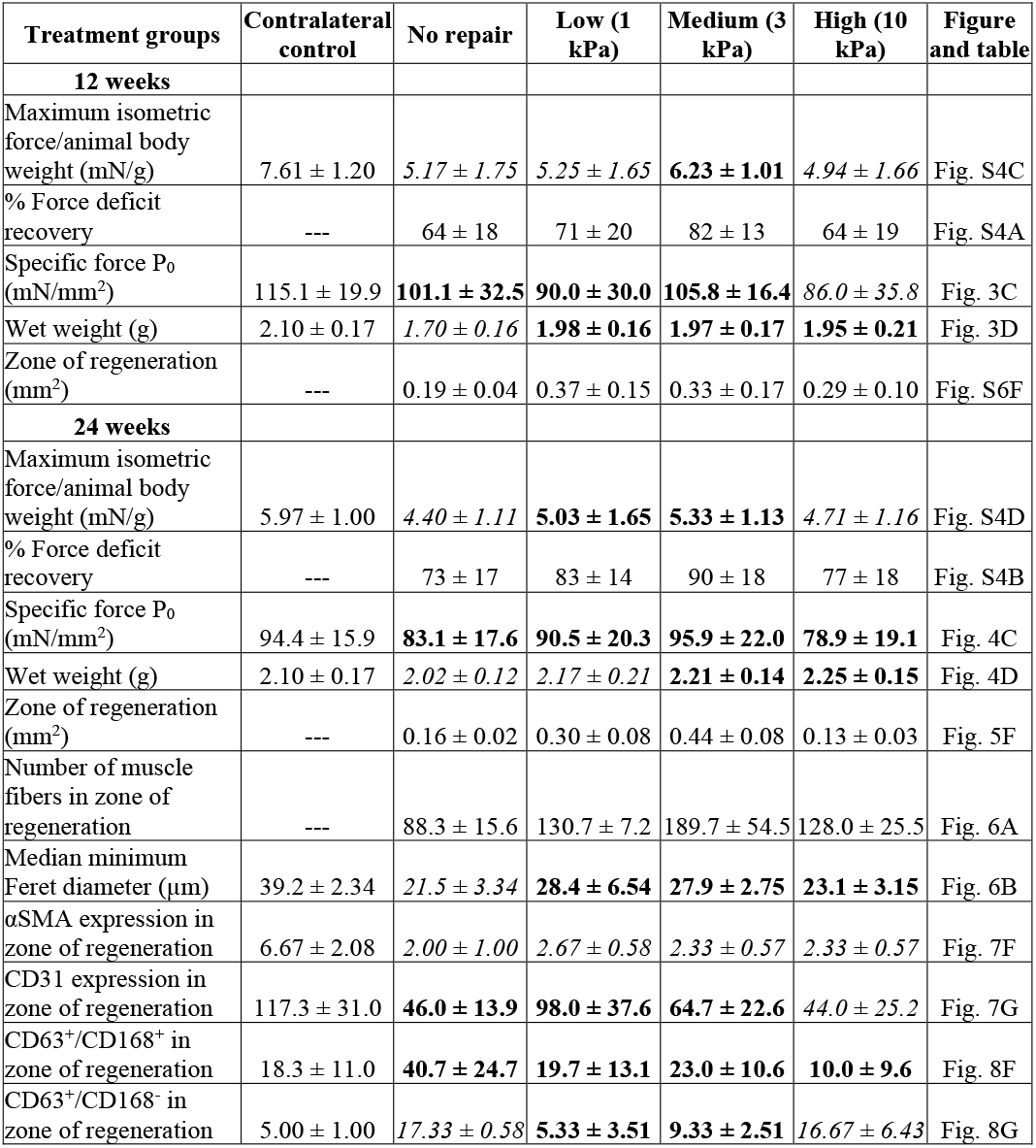
Summary of experimental findings across treatment groups at both 12 and 24 weeks post-VML.

Values are presented as mean ± standard deviation expect for the zone of regeneration at 12 weeks and median minimum Feret diameter which are reported as the median and 1^st^ and 3^rd^ quartiles. Values denoted with the bolded lettering were not significantly different (*P* > 0.05) from the contralateral control, while values with italic lettering were significantly different (*P* < 0.05) from the contralateral control muscles. SMA, smooth muscle actin; ---, analysis not applicable to contralateral control.

While our results showed modest enhancement of functional recovery following VML, they fall short of reaching native muscle properties. A recent meta-analysis exploring regenerative therapies following VML suggests that currently the most effective treatments for functional regeneration of skeletal muscle combine biomaterials seeded with cells^67^. Previous work by our lab supports these findings where a decellularized porcine bladder extracellular matrix (BAM) reseeded with myogenic cells showed substantial functional recovery of a rat LD muscle 24 weeks post-injury^3^. Moreover, this tissue engineered muscle repair construct (TEMR) has been shown to augment skeletal muscle recovery post-VML in a number of different animal models^44,68,69^. This enhancement in functional outcomes is likely linked to the presence of a cellular component that can rapidly fill the necrotic core of the injury, modulate the immune response by release of cytokines, and facilitate myogenesis^49,55,67^. However, in order to elucidate the specific role material biophysical properties play in muscle repair, we chose to implant hydrogels without encapsulated cells in this study. While this material system was unable to elicit complete restoration of muscle function, the findings of this study should inform subsequent biomaterial design for the repair of complex injuries.

Following functional testing, histological staining was used to further examine the mechanisms governing VML repair as a function of hydrogel elasticity. Masson’s trichrome staining revealed a zone of regenerating muscle characterized by small, disorganized muscle fibers, neighboring uninjured native muscle at the material-tissue interface (**Fig. 5, S6**). Quantification of this zone of regeneration showed that hydrogel-treated tissues contained larger regenerative areas than non-treated muscles at 12 weeks post-injury; however, these results were not statistically significant. Analysis at 24 weeks revealed a significantly larger zone of regeneration in medium stiffness hydrogel treated muscles compared to other experimental groups. These findings corroborate our functional testing results suggesting that superior myogenesis is the driver of improved functional outcomes. However, it is important to note that despite evidence of myogenesis at the material-tissue interface a significant amount of fibrotic tissue remained at the center of the defect. Previous work has described that the inclusion of myogenic cells with a regenerative therapeutic is likely necessary for complete restoration of muscle function and volume following VML^3,44,67^. Therefore, future studies may look to assess skeletal muscle regeneration following implantation of cell-laden HA hydrogels.

To quantitatively evaluate the extent of myogenesis, muscle minimum Feret diameter and fiber cross sectional area (FCSA) were evaluated in the zone of regeneration and compared to native muscle. Our analysis revealed that muscles treated with medium stiffness hydrogels supported significantly more *de novo* myogenesis than other experimental groups (**Fig. 6**). We believe these superior regenerative outcomes are due to a balance of improved cell infiltration compared to high stiffness hydrogels as well as mechanotransduction-mediated signaling to facilitate myogenic growth and differentiation. Additionally, analysis of muscle minimum Feret diameter showed that hydrogel treated muscles contained median fiber sizes that were statistically similar to native muscle, while NR tissues showed significantly reduced fiber sizes. These results corroborate our functional testing data and histological analysis in the zone of regeneration, underscoring the important role material mechanical properties can play in cell response and wound healing post-VML. Despite this improvement, there remains a marked increase in the frequency of small diameter muscle fibers at the zone of regeneration regardless of treatment, which is indicative of immature muscle. As a result, there remains ample room for therapeutic advancement to aid in myogenic recovery, perhaps by including of cells or other bioactive moieties.

Angiogenesis and vascularization of skeletal muscle following VML is essential for sustained functional recovery due to the high metabolic demands of muscle tissue^13^. IHC staining for pericytes (αSMA) was significantly reduced in all experimental groups indicating that hydrogel intervention did not facilitate the development of mature vascular structures such as arteries and veins (**Fig. 7**). However, lower stiffness hydrogels (low: 1 kPa; medium: 3 kPa) supported levels of CD31^+^ cells that were not statistically different from contralateral control muscles. This result could be attributed to quicker cell migration into hydrogels with fewer crosslinks as well as the inherent proangiogenic characteristics of HA^54,56^. However, these results clearly indicate that the extent of revascularization following VML still needs to be improved. Recent work by our lab and others has shown that incorporation of heparin into hydrogels can sequester soluble growth factors and greatly improve angiogenic outcomes^15,70,71^. As a result, future experiments could look to augment revascularization through the addition of bioactive components to the hydrogel system.

It is important to note that macrophages are typically activated within hours of skeletal muscle injury and support the inflammatory and repair processes for roughly 14 days before returning to baseline levels^49,50^. During the later phase of wound healing a transition from an initial inflammatory response to a type 2 immune response is characterized by a change in the macrophage polarization from the classically activated M1 phenotype to the alternatively activated M2 phenotype^49–51^. This temporal change is linked to a shift from proliferation of myogenic cells to differentiation and ultimately myogenesis^49,50,72^. However, VML injuries are characterized by an aberrant wound response where persistent inflammation limits regenerative outcomes^31^. As a result, while evaluating the presence of macrophages at 24 weeks post-VML cannot be used to assess the initial inflammatory response, understanding sustained macrophage activity and polarization post-VML is critical for skeletal muscle repair. While no statistical differences were observed in M2 macrophage staining (CD68^+^/CD163^+^), M1 macrophages (CD68^+^/CD163^-^) remained significantly elevated in NR and high stiffness hydrogel treated muscles compared to native muscle, suggesting a chronic inflammatory response (**Fig. 8**). In contrast, levels of both M1 and M2 macrophages in muscles treated with lower stiffness hydrogels were similar to native muscle. Interestingly, this result correlates strongly to the limited amount of *de novo* myogenesis detected at the zone of regeneration.

## 5. Conclusions

This study aimed to inform biomaterial design for the treatment of VML injuries by elucidating the role that hydrogel mechanical properties play in skeletal muscle wound healing and functional recovery. Prior work has shown that myogenic cell growth and differentiation can be influenced by substrate elasticity via mechanotransduction *in vitro*; however, it remains unclear if these findings directly translate to tissue engineering strategies *in vivo*. In the present study, we developed a photoreactive hydrogel platform with customizable mechanical properties for repair of VML injuries. Light-mediated thiol-ene click chemistry allowed for *in situ* hydrogel polymerization, spanning a range of physiologically relevant mechanical properties, in a rat LD model of VML. This gelation scheme allows for facile hydrogel delivery that could be readily administered to the defect site and conform to injuries of variable sizes and geometries. NorHA hydrogels with stiffnesses approximating developing muscle (medium: 3 kPa), as opposed to mature (high: 10 kPa) tissue, supported improved functional recovery and myogenesis while limiting fibrosis at 12 weeks post-VML. Moreover, the magnitude of functional recovery was maintained at 24 weeks post-injury, indicating that the extent of repair was durable over time. Histological analysis of the defect-muscle interface also revealed that muscles treated with medium stiffness hydrogels contained the largest zone of regeneration and *de novo* myogenesis. These improved regenerative outcomes were substantiated by an increase in the number of muscle fibers at the zone of regeneration when compared to non-treated muscles and a median minimum Feret diameter that was not statistically different from native muscle. Finally, lower stiffness hydrogels displayed similar levels of macrophage infiltration and polarization to native muscle, attenuating the significantly increased M1 macrophage levels observed in non-repaired or high stiffness hydrogel treated muscles. While there remains significant room for therapeutic optimization, our findings illustrate that MMP-degradable hydrogels with stiffness that may mimic developmental muscle tissue can facilitate improved functional recovery and endogenous muscle repair following VML injury.

## Supporting Information

^1^H NMR spectra for NorHA, MALDI spectra for the dithiol peptide, and additional data from the VML animal model results can be found in the Supporting Information.

## Acknowledgments

The authors would like to acknowledge the University of Virginia’s Center for Advanced Biomanufacturing for use of its rheometer, Dr. Rachel Letteri for use of her peptide synthesizer, the University of Virginia’s Biomolecular Analysis Facility for MALDI analysis, the University of Virginia’s Rapid Prototyping Lab for printing hydrogel molds, Areli Rodriguez for assistance with rat functional testing, and Sarah Dyer for help with histological and immunohistochemical staining. This work was supported by seed funding from the University of Virginia’s Center for Advanced Biomanufacturing, the UVA Biotechnology Training Program (T32GM136615), and the NIH (R21AR075181, R01AR078886). The content is solely the responsibility of the authors and does not necessarily represent the official views of the National Institutes of Health.

