## Supplementary Information for "Photoreactive hydrogel stiffness influences volumetric muscle loss repair"

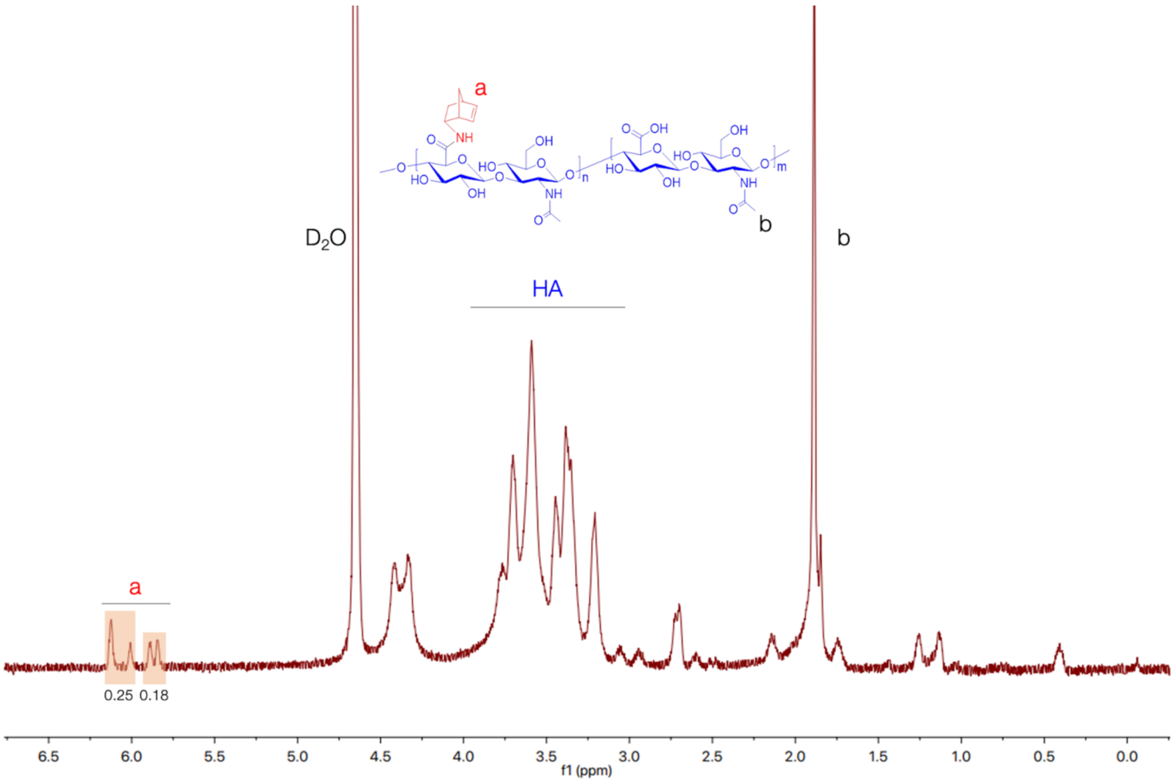


**Fig. S1: $^1$H NMR spectrum of norbornene-modified hyaluronic acid (NorHA).** Spectrum indicates ~ 22% modification of HA repeat units with norbornene groups.

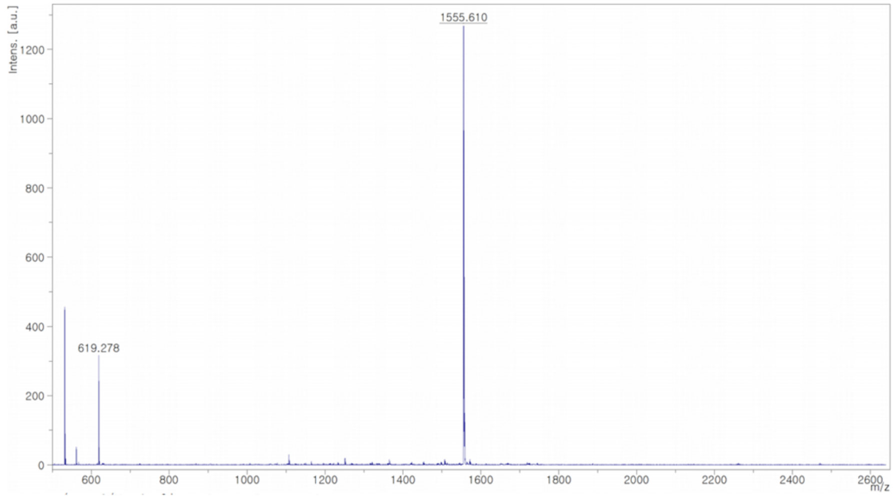


**Fig. S2: MALDI spectrum of MMP-degradable dithiol crosslinker.** Degradable dithiol peptide with sequence GCNSVPMSMRGGSNCG. Expected mass: 1557 g/mol. Actual mass: 1555 g/mol.

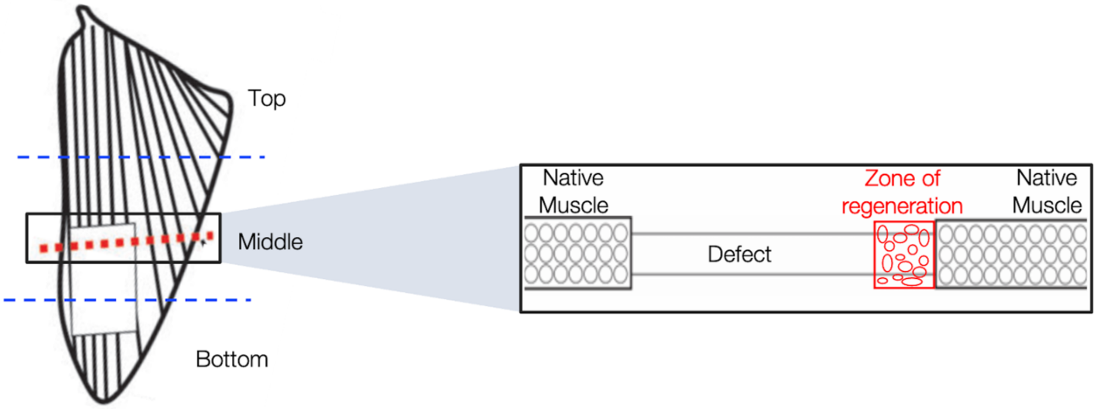


**Fig. S3: Schematic representation of zone of regeneration for histological and immunofluorescent imaging.** Cross-sectional images were acquired as confocal tile scans. The zone of regeneration was characterized by disorganized smaller diameter muscle fibers. The VML defect was marked by a thin layer of connective tissue with minimal cell infiltration. Schematic created with BioRender.

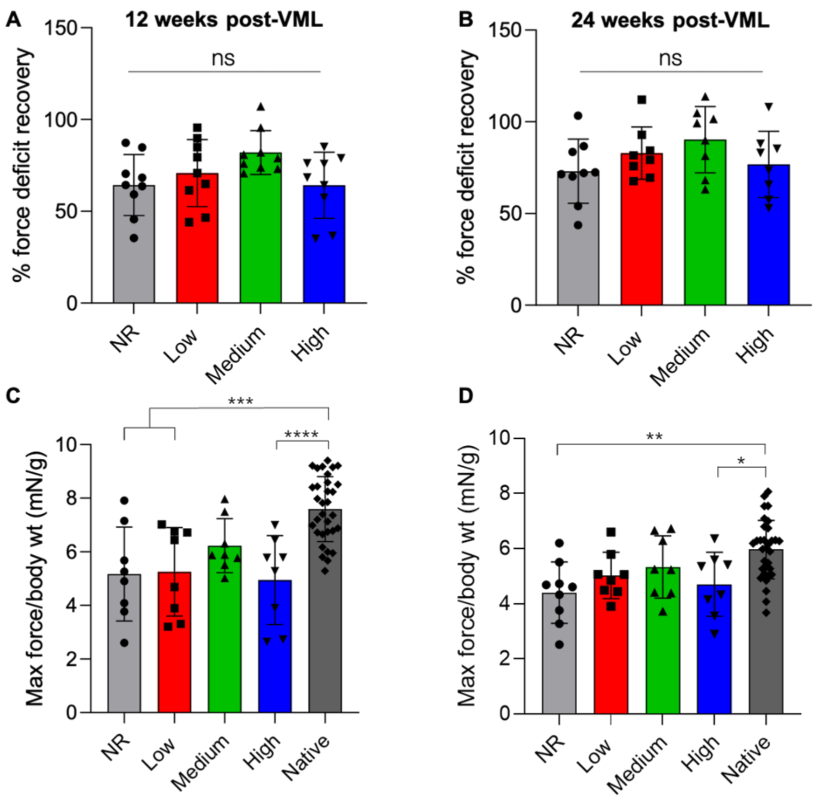


**Fig. S4: *Ex vivo* % force deficit recovery and maximum isometric force normalized to animal body weight.** *Ex vivo* maximum force of contraction was normalized to the maximum force produced by contralateral control muscles. There were no statistical differences between % force deficit recovery at (A) 12 weeks or (B) 24 weeks. (C) Maximum isometric contraction force normalized to animal body weight indicated that medium stiffness hydrogels supported muscle force generation that was not statistically different from the native LD at 12 weeks. (D) Similarly, at 24 weeks low and medium stiffness hydrogels supported force generation that was not statistically different from the native LD.* $P < 0.05$, ** $P < 0.01$, *** $P < 0.001$, **** $P < 0.0001$. Data presented as mean +/- SD.

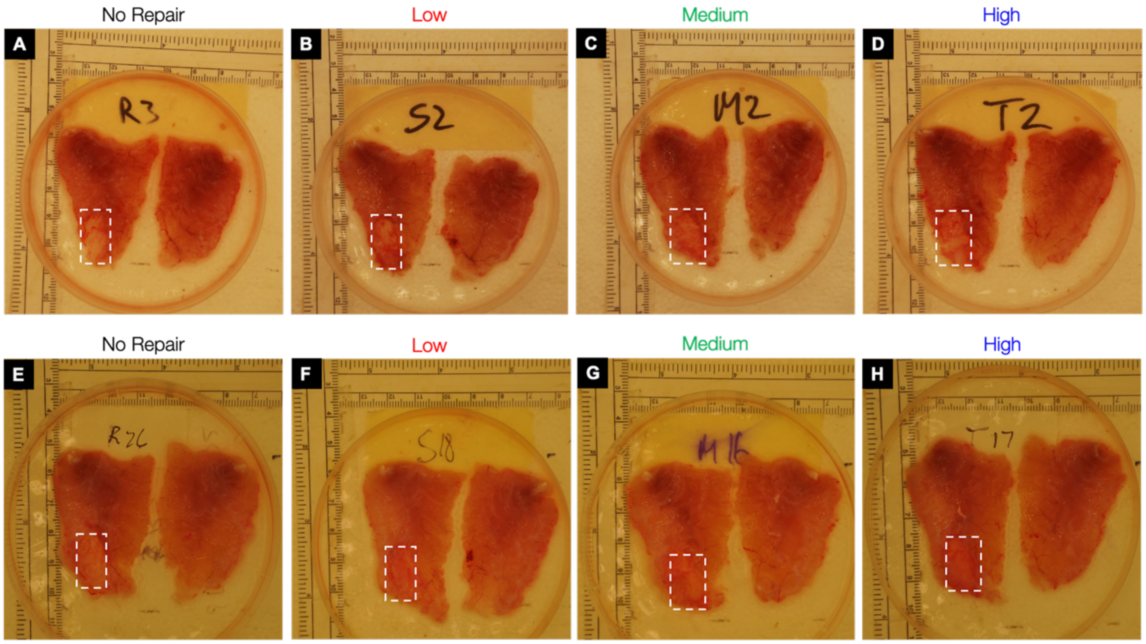


**Fig. S5: Gross tissue morphology 12 and 24 weeks post-VML.** At 12 weeks post-implantation, gross tissue morphology suggests superior tissue regeneration at the VML injury site (*white squares*) in hydrogel groups compared to no repair injuries. Similarly, after 24 weeks hydrogel treated tissues appear to have increased muscle volume at the defect site (*white squares*).

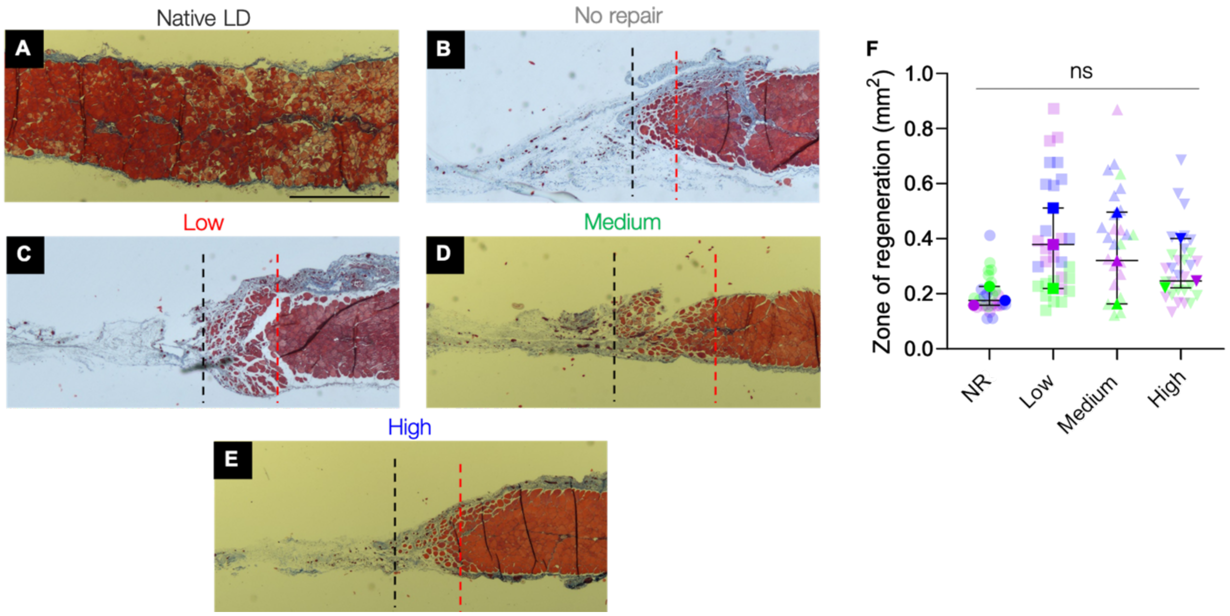


**Fig. S6: Histological analysis of LD muscles at 12 weeks.** (A-E) Tissue and cell morphology at the interface between native muscle and the VML injury was visualized by Masson's trichrome staining (muscle tissue in red, collagen deposition in blue, and nuclei in black). The area between the dashed red and black lines represents the zone of regeneration characterized by small disorganized myofibers. The red line denotes the interface between native muscle and VML injury site and the black line represents the last point at which detectable myofibers were present. (F) Quantification of the zone of regeneration at 12 weeks post-injury indicated similar levels of myogenesis across treated muscles. Colors denote different experimental muscles (biological replicates, $n$ = 3 per group) with solid shapes representing the median and translucent shapes representing individual muscle sections (technical replicates, $n$ = 10). Data presented as the median across 10 muscle sections. Scale bar: 1 mm.

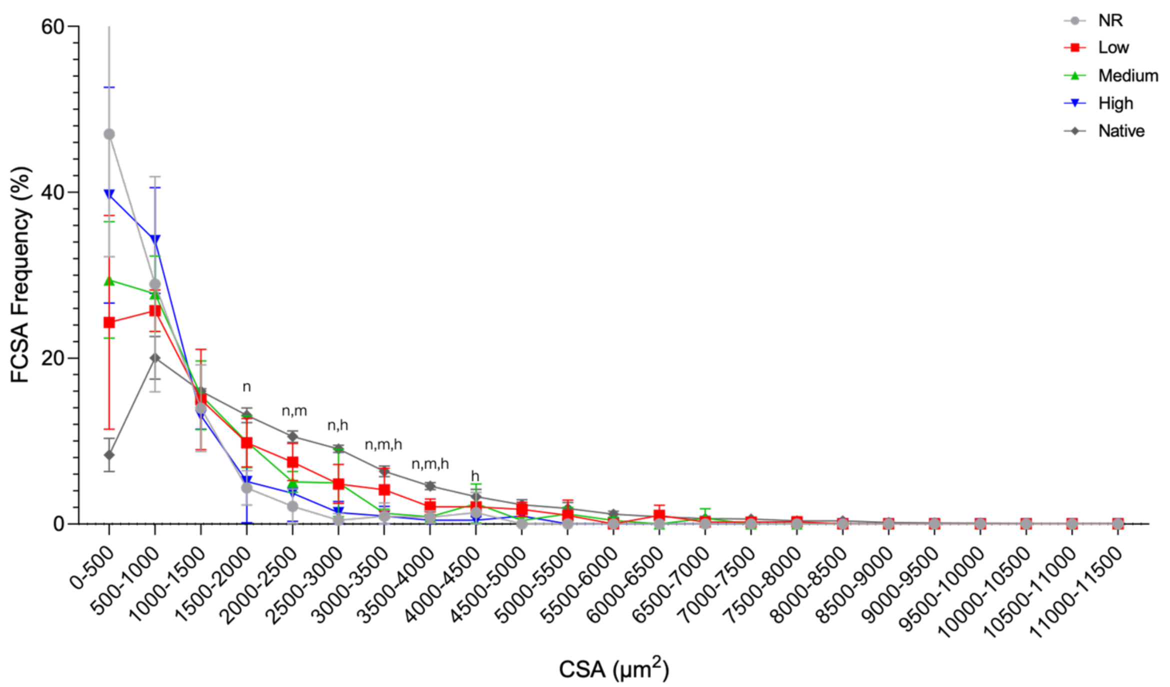


**Fig. S7: Muscle fiber cross sectional area (FCSA) analysis at 24 weeks post-VML.** FCSA frequency distribution curve showed a leftward shift toward smaller muscle fibers compared to uninjured muscle regardless of treatment type. Statistically significant differences from native muscle are denoted by *n* (NR), *l* (low), *m* (medium), and *h* (high). Data presented as mean +/- SD. *n* = 3 muscles per experimental group.

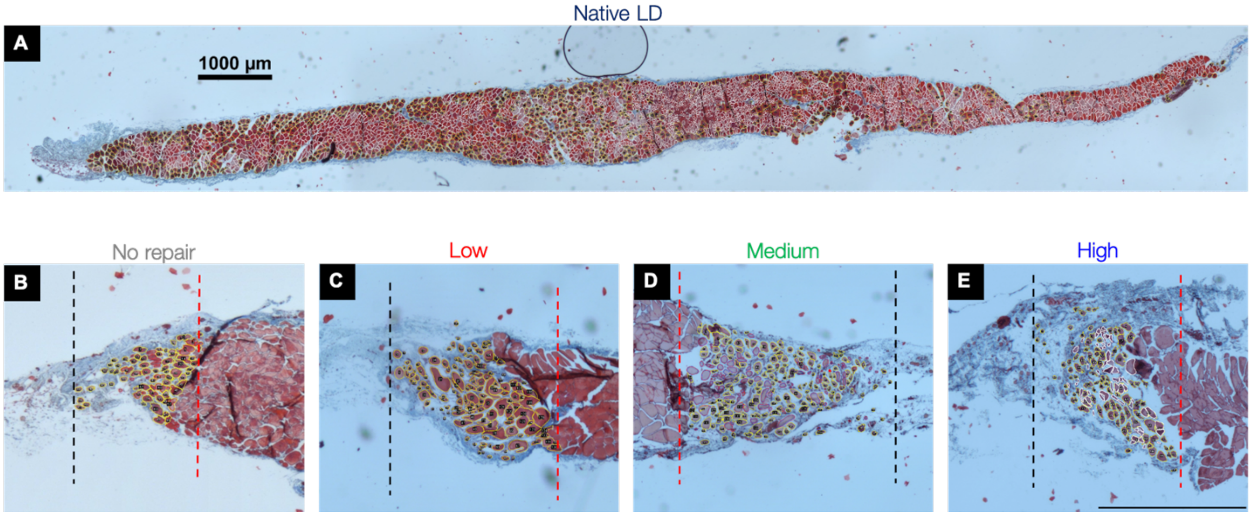


**Fig. S8: Analysis of minimum Feret diameter at 24 weeks post-VML.** (A) Minimum Feret diameter was quantified throughout the entirety of the LD muscle section using a semi-automated ImageJ processing pipeline and manual counting. (B-E) Minimum Feret diameter of regenerating fibers was assessed at the zone of regeneration across each experimental group using the same ImageJ pipeline and manual counting. *n* = 3 muscles per experimental group. Scale bars: 1000 µm.
